# Reduced cellular diversity and an altered basal progenitor cell state inform epithelial barrier dysfunction in human type 2 immunity

**DOI:** 10.1101/218958

**Authors:** Jose Ordovas-Montanes, Daniel F. Dwyer, Sarah K. Nyquist, Kathleen M. Buchheit, Chaarushena Deb, Marc H. Wadsworth, Travis K. Hughes, Samuel W. Kazer, Eri Yoshimoto, Neil Bhattacharyya, Howard R. Katz, Tanya M. Laidlaw, Joshua A. Boyce, Nora A. Barrett, Alex K. Shalek

## Abstract

Tissue barrier dysfunction is a poorly defined feature hypothesized to drive chronic human inflammatory disease^1,2^. The epithelium of the upper respiratory tract represents one such barrier, responsible for separating inhaled agents, such as pathogens and allergens, from the underlying submucosa. Specialized epithelial subsets—including secretory, glandular, and ciliated cells—differentiate from basal progenitors to collectively realize this role^3-5^. Allergic inflammation in the upper airway barrier can develop from persistent activation of Type 2 immunity (T2I), resulting in the disease spectrum known as chronic rhinosinusitis (CRS), ranging from rhinitis to severe nasal polyps^6-8^. Whether recently identified epithelial progenitor subsets, and their differentiation trajectory, contribute to the clinical presentation and barrier dysfunction in T2I-mediated disease in humans remains unexplored^3,9,10^. Profiling twelve primary human samples spanning the range of clinical severity with the Seq-Well platform^11^ for massively-parallel single-cell RNA-sequencing (scRNA-seq), we report the first single-cell transcriptomes for human respiratory epithelial cell subsets, immune cells, and parenchymal cells (18,036 total cells) from a T2I inflammatory disease, and map key mediators. We find striking differences between non-polyp and polyp tissues within the epithelial compartments of human T2I cellular ecosystems. More specifically, across 10,383 epithelial cells, we identify a global reduction in epithelial diversity in polyps characterized by basal cell hyperplasia, a concomitant decrease in glandular and ciliated cells, and phenotypic shifts in secretory cell function. We validate these findings through flow cytometry, histology, and bulk tissue RNA-seq of an independent cohort. Furthermore, we detect an aberrant basal progenitor differentiation trajectory in polyps, and uncover cell-intrinsic and extrinsic factors that may lock polyp basal cells into an uncommitted state. Overall, our data define severe T2I barrier dysfunction as a reduction in epithelial diversity, characterized by profound functional shifts stemming from basal cell defects, and nominate a cellular mechanism for the persistence and chronicity of severe human respiratory disease.

## MAIN TEXT

T2I plays a dual role in regulating homeostatic processes, such as metabolism^12^, and promoting inflammatory defense mechanisms in response to parasites, venoms, allergens, and toxins^13^. T2I is characterized by sensor cells (epithelia, macrophages, dendritic cells, mast cells) producing first-order cytokines (TSLP, IL-25, IL-33) that cause release of second-order cytokines (IL-4, IL-5, IL-13, AREG) from lymphocytes^5,14^. This, in turn, helps recruit effector cells (eosinophils, basophils, mast cells, monocytes), and induces epithelial remodeling, such as goblet cell hyperplasia^15^, to restore tissue integrity^5,14^. Accumulating evidence suggests that immune modules, which are productively activated in acute settings to restore homeostasis, may reach an aberrant set-point during chronic inflammatory diseases such as CRS^16,17^. For example, cytokine modules may become self-reinforcing in T2I leading to substantial alterations in gross tissue architecture^18^ and the reduced epithelial barrier integrity characteristic of severe clinical presentations, such as polyps^19,20^. Whether and how human cellular tissue ecosystems shift in composition and function during chronic T2I disease remains unknown.

To directly address this in respiratory barrier tissue, we used Seq-Well^11^ (massively-parallel scRNA-seq for unique molecular identifier (UMI)-collapsed digital gene expression profiling) to profile surgically resected and dissociated sinus tissue from patients across a spectrum of CRS severity (**Fig. 1a; Supplementary Table 1; Methods; n=12 samples)**. Deconstructing these tissues into their component cells provides a unique lens into the ecosystem of human T2I, allowing us to: 1. characterize each major cell type absent the biases typically introduced by flow sorting/marker selection; 2. reconstruct tissue-level interactions; and, 3. rapidly assess which cell types/states have striking disease-associated transcriptional differences. To the latter point, our scRNA-seq data spans tissue specimens ranging from mild chronic inflammation and no eosinophilia, through moderate eosinophilia, to more severe eosinophilic rhinosinusitis, presenting an opportunity to identify cellular features that correlate with T2I inflammatory disease severity and gross anatomical features, such as the presence of polyps (n_Non-Polyp_=6; n_Polyp_=6; **Supplementary Table 1**)^8^. As no healthy individuals were clinically indicated for nasal tissue resections during enrollment (**Methods**), and nasal brushings cannot fully recapitulate the nasal mucosal ecosystem given a bias towards apical cells^21^, we focused on globally assessing the cell types/states that inform the spectrum of T2I-mediated CRS **(Fig. 1)** and its gross anatomical manifestations (e.g., polyps vs non-polyps)^5,8,19,20^.

**Figure 1.**
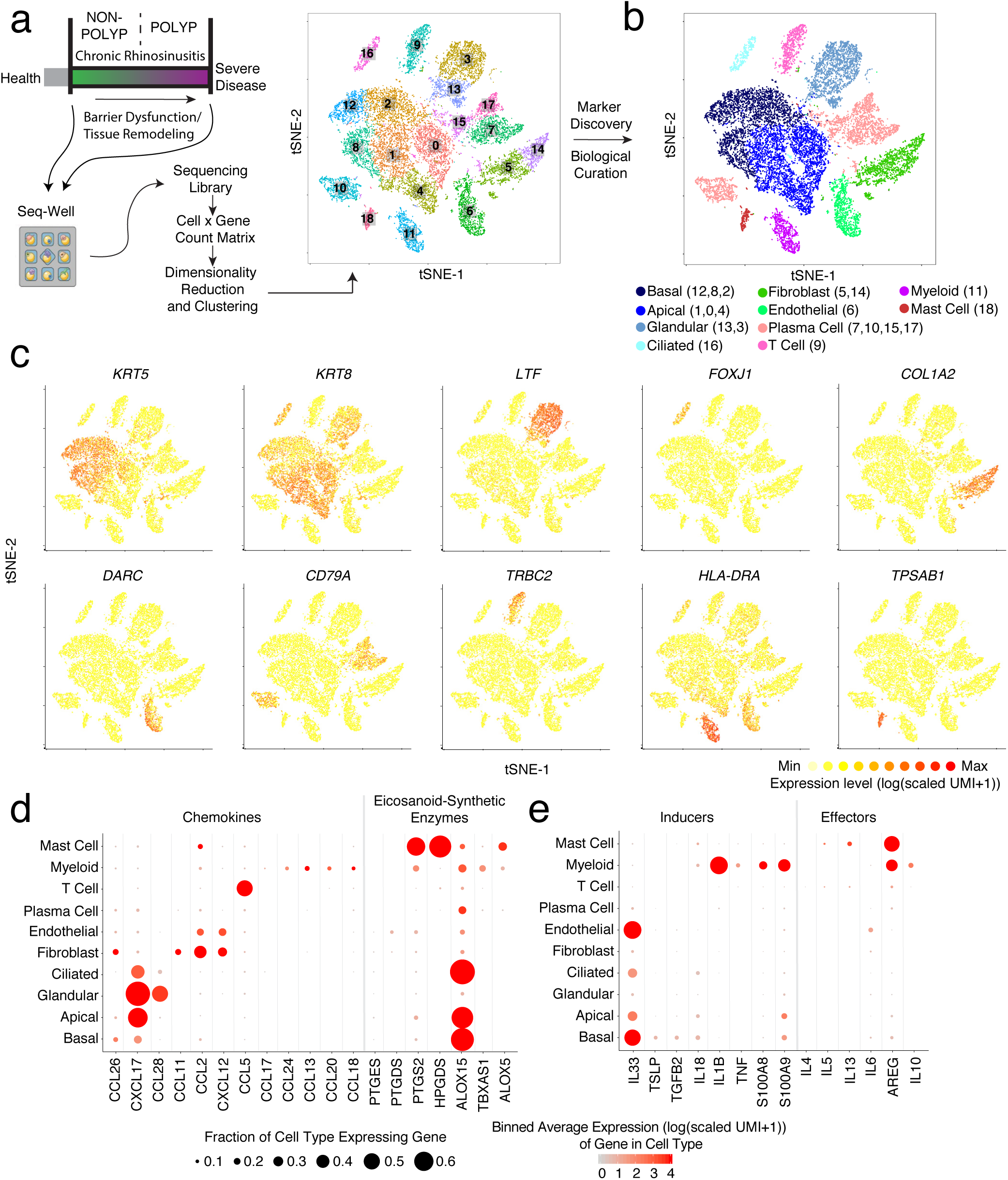
The inflamed human nasal cellular ecosystem by scRNA-seq provides a map for type 2 immunity mediators. **(a)** Illustration of the clinical disease spectrum sampled (n=12 samples) and the experimental workflow leading to the generation of a tSNE plot displaying 18,036 single cells, colored by clusters identified through shared nearest neighbor (SNN) analysis (**Supplementary Table 3; Methods**) from respiratory tissue. **(b)** tSNE plot of 18,036 single cells (n=12 samples), colored by cell types identified through marker discovery (ROC test) and biological curation of identified clusters (**Supplementary Table 3; Methods**). **(c)** Select marker gene overlays displaying binned count-based (unique molecular identifier (UMI) collapsed) expression level (log(scaled UMI+1) on a tSNE plot from (**b**) for key cell types identified (see **Supplementary Table 3** for full gene lists); area under the curve (AUC) 0.998 to 0.7 for all markers displayed. **(d)** Dot plot of chemokines and lipid mediators with known roles in type 2 immunity mapped onto cell types across all samples, dot size represents fraction of cells within that type expressing, and color intensity binned count-based expression level (log(scaled UMI+1)) amongst expressing cells (see **Supplementary Figure 4** by disease state). **(e)** Dot plot of inducers and effectors of type 2 immunity mapped onto cell types across all samples, dot size represents fraction of cells within that type expressing, and color intensity binned count-based expression level (log(scaled UMI+1)) amongst expressing cells (see **Supplementary Figure 4** by disease state).

After library preparation and sequencing, we derived a unified cells-by-genes expression matrix containing digital gene expression values for all cells passing quality thresholds (n=18,036**; Supplementary Fig. 1a,b; Supplementary Table 2; Methods)**. We then performed dimensionality reduction and graph-based clustering^22^ **(Fig. 1a; Methods)**. Identified clusters were present across all patients within the non-polyp and polyp groups, with no cluster composed solely of cells from a single individual **(Supplementary Fig. 1c,d; Supplementary Table 3)**. We subsequently derived lists of cluster-specific genes **(Supplementary Table 3),** and used them to define our parenchymal and immune cell subsets^22^ **(Fig. 1b)**. From the complete lists of cluster-specific/sensitive markers, we display several lineage-defining genes to highlight the major cell types recovered, including basal (*KRT5*), apical (*KRT8*), ciliated (*FOXJ1*) and glandular (*LTF*) epithelium, and associated endothelial (*DARC*), fibroblast (*COL1A2*), plasma (*CD79A*), myeloid (*HLA-DRA*), T (*TRBC2*), and mast cells (*TPSAB1*) **(Fig. 1c).**

Despite the potential challenges associated with isolating single parenchymal and immune cells from tissues^23^, we sequenced and annotated a highly reproducible distribution of cell types across patients **(Supplementary Fig. 1c,d)**. Collectively, we recovered abundant cell numbers for all expected cell types^5,6,8,18^, save for ILC2s and eosinophils **(Supplementary Fig. 1d)**. Given the rarity of ILCs in unfractionated tissue^24^, we are likely underpowered to recover a distinct cluster **(Methods)**. Meanwhile, when examining our data for *CLC* expression, we did find a small set of eosinophils within our myeloid cluster **(Supplementary Fig. 2a).** Nevertheless, their frequency was much lower than anticipated based on both the literature^8^ and matched clinical pathology reports. Since we measured a robust eosinophil and Th2 signature in snap-frozen whole tissue nasal RNA-Seq libraries from nasal polyps^8^ **(Supplementary Fig. 2b,c,d)**, and they can be recovered by flow cytometry, our data suggest that eosinophil transcripts may be particularly susceptible to degradation during isolation, likely because several of their lineage-defining genes include RNAses^25^ **(Methods)**.

Lymphoid and myeloid cells are recruited and positioned in tissues to facilitate effector functions during T2I through the action of chemokines and lipid mediators, but the cell of origin for these molecules can be obscured by bulk cellular analyses^6,26^. The eotaxins (*CCL26, CCL11, CCL24*) are chemotactic for eosinophils, acting through CCR3 (*CCL2, CCL5* and *CCL13* are also CCR3 ligands)^25^. In our scRNA-Seq data, we detected each eotaxin in distinct sources, including basal cells (*CCL26*), fibroblasts (*CCL26* and *CCL11*), and myeloid cells (*CCL24*) **(Fig. 1d)**. The mucosal cytokine *CXCL17*^27^ was largely detected in apical and glandular cells, while *CCL28*, a chemokine involved in the positioning of IgA+ plasma cells in mucosal tissues^28^, was specifically detected in glandular epithelium **(Fig. 1d)**. Lipid mediators, such as prostaglandins, thromboxane, and leukotrienes, also play key roles in T2I^29^. We found mast cells specifically enriched for *HPGDS* and *PTGS2*, and *ALOX5,* suggesting they may be a dominant source of prostaglandin D2 and prostaglandin E2, implicated in intestinal epithelial repair^30^, and leukotriene biosynthesis within CRS, respectively. Myeloid cells, meanwhile, enriched for *TBXAS1*, responsible for thromboxane A2 biosynthesis, and basal, apical and ciliated epithelial cells were high expressers of *ALOX15*, recently reported to regulate ferroptotic cell death in airway epithelial cells^31^ **(Fig. 1d)**.

The production of instructive first-order cytokines, alongside these chemokines and lipid mediators, primes recruitment and activation of effector mechanisms. In particular, IL-25, IL-33^32-34^, and TSLP^35,36^ are broadly referred to as epithelial-derived cytokines^5,6,26^, and yet, beyond a study illustrating IL-33 production by basal cells in human COPD^37^, little is known about their cell-of-origin in human disease. Consistent with previous reports, we did not detect *IL25* in our system^38^ **(Fig. 1e)**. *IL33* was present in both basal and endothelial cells, and to a lesser extent apical and ciliated cells, and *TSLP* restricted to basal cells **(Fig. 1e)**. Furthermore, amplifying cytokines (e.g. IL-1 family members)^5^ were either distributed throughout both epithelial and immune subsets (*IL18*), or limited to myeloid cells (*IL1B*).

With regard to second-order cytokines produced in response to TSLP and IL-33, we detected a small, but consistent, subset of CD4+ T cells expressing *IL4*, *IL5*, *IL13, and HPGDS*, fitting the recently described profile of allergen-specific Th2A cells^39^ **(Supplementary Fig. 3a; Methods).** All identified T cell sub-clusters scored evenly for a set of TCR complex genes **(Supplementary Fig. 3b; Methods),** and we could not detect robust signals for Type-1 or Type-17 inducer or effector cytokines across any cell subset **(Supplementary Fig. 3c).** Beyond T cells, we also noted a substantial number of mast cells producing *IL5* and *IL13* **(Fig. 1e)^35^**. Furthermore, mast, along with myeloid, cells were the main expressers of the tissue-reparative cytokine *AREG^40^* **(Fig. 1e)**. Patients with or without gross structural differences (i.e., polyps) showed consistent cells-of-origin for the T2I-related chemokines, lipids, and cytokines, except for select mediators, such as *CCL26* and *AREG*, which were more broadly detected in polyp tissue, and *PTGS2* (e.g. COX2), which was reduced in polyp epithelium **(Supplementary Fig. 4a,b)**. Overall, our data highlight the activation of similar T2I modules across CRS tissues, implicate basal cells as key producers of first-order cytokines, and CD4+ T cells and mast cells of second-order cytokines, with the potential to influence epithelial cell states^43,44^.

To better understand the epithelial cell states present across CRS, we further analyzed clusters within our broad epithelial cell populations **(Fig. 2a; Supplementary Fig. 5a,b,c).** We provide, to our knowledge, the first single-cell human transcriptomes in the absence of flow-sorting for basal^45^, secretory, glandular, and ciliated cell types from a T2I ecosystem **(Fig.2a,b,c; Supplementary Fig. 5; Supplementary Table 3)**. Marker gene analysis across epithelial cell types (e.g. basal vs. differentiating/secretory vs. glandular vs. ciliated) identified conserved programs present in the three clusters with a basal phenotype (*TP63, KRT5,* and high basal cell score^10^), the three with a differentiating/secretory phenotype (*KRT8, SERPINB3, SCGB1A1,* and baseline basal cell score^10^), the two with a glandular phenotype (*LTF, TCN1, LYZ*), and the one with a ciliated phenotype (*CAPS, OMG, FOXJ1, PIFO*) **(Fig. 2a,b; Supplementary Fig. 5a,b,d; Supplementary Table 3)**^3,4^. Marker gene analysis within each epithelial cell type (e.g. 12 vs. 8 vs. 2 only), meanwhile, revealed further granularity associated with the presence or absence of polyps **(Fig. 2c,d; Supplementary Table 3)**. Within basal cells and differentiating/secretory cells, we observed that canonical Type 2 cytokine induced genes (*POSTN^46^* and *ALOX15*) amongst others (**Supplementary Table 3**), drive clustering, suggesting that the separations we observe may be mediated by differences in the sensing or impact of cytokines between disease states **(Fig. 2c,d)**. Importantly, a canonical correlation analysis (CCA)^47^, performed across non-polyp and polyp samples, returned similar basal and differentiating/secretory cluster groupings **(Supplementary Fig. 5c).**

**Figure 2.**
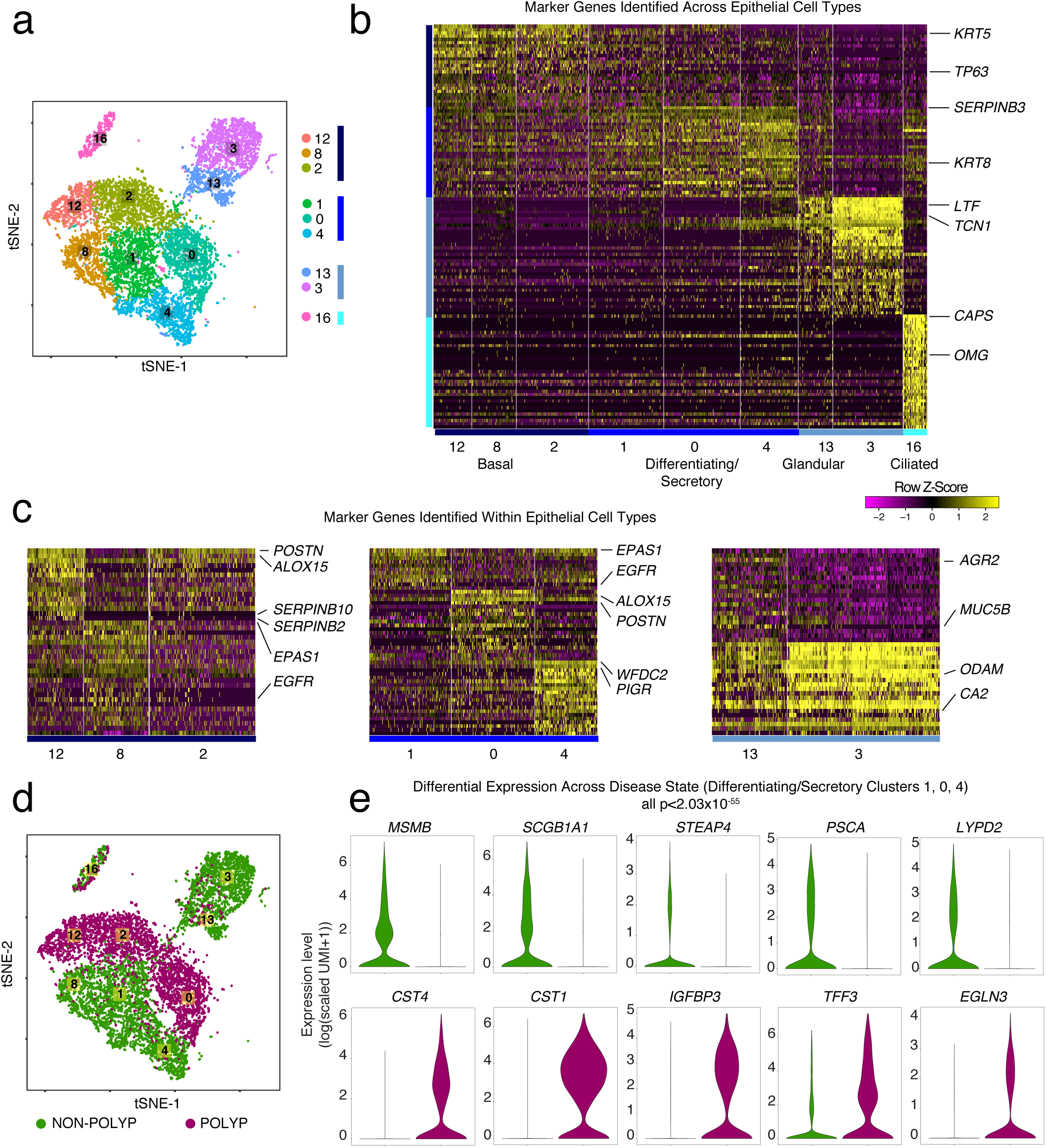
Single-cell transcriptomes of epithelial cells in type 2 inflammation and stratification by disease state. **(a)** tSNE plot of 10,274 epithelial cells (n=12 samples), colored by clusters identified through SNN, with adjacent color bars representing related cell clusters based on marker genes from ROC-tests (AUC > 0.65), hierarchical clustering, and scoring over basal cells (Cluster Numbers as in **Fig. 1a,b**; see **Methods** and **Supplementary Fig. 5**). **(b)** Row-normalized heatmap of the top marker genes identified by ROC-test (AUC > 0.65) for each epithelial cell type with select genes displayed on y-axis and cluster annotations on x-axis (see **Supplementary Table 3** for full gene lists). **(c)** Row-normalized heatmap of the top marker genes identified by ROC-test (AUC > 0.6) within each cell type for each cell cluster with genes displayed on y-axis and cluster annotations on x-axis (see **Supplementary Table 3** for full gene lists). **(d)** tSNE plot of 10,274 epithelial cells colored by disease state (Cluster Numbers as in **Fig. 1a,b**; n= 6 non-polyp, n=6 polyp samples). **(e)** Violin plots for the count-based expression level (log(scaled UMI+1)) for key differentially expressed genes using bimodal test within the differentiating/secretory cell subset across disease states; n=6 non-polyp, n=6 polyp samples, *bimodal test, all p<2.03x10^−55^ or less with Bonferroni correction for multiple hypothesis testing based on number of genes tested.

The two predominant secreted mucins of the upper respiratory tract are MUC5B and MUC5AC, which differ in their biophysical properties and potential to mediate microbe adhesion^48^. Although the production of both has been commonly ascribed to a set of goblet cells in human airway^4^, we observed that expression of *MUC5B* is restricted to glandular cells (cluster 13) in nasal mucosa, which do not express *MUC5AC* (r=-0.0375; p=0.4952, corrected Holm’s method**; Fig. 2c; Supplementary Fig. 5e)**. Instead, *MUC5AC* is expressed in a subset of secretory cells (clusters 0 and 4) co-expressing *SCGB1A1* and *FOXA3* (*MUC5B* vs *AZGP1*: r=0.491, *MUC5AC* vs *AZGP1*: r=-0.084, *MUC5AC* vs *SCGB1A1*: r=0.184, *MUC5AC* vs *FOXA3*: r=0.181; p<0.0001, corrected Holm’s method**; Supplementary Fig. 5e,f)**. This suggests that the goblet cell program is layered atop a secretory cell base^49-53^, and that glandular cells are the predominant source of *MUC5B*^54^. Expression of *SPDEF*, a putative goblet cell transcription factor^4^, was shared amongst mucin producing cells, but relatively enriched in *MUC5AC* expressing cells (SPDEF vs MUC5B: r=0.107, SPDEF vs MUC5AC: r=0.213; p<0.0001, corrected Holm’s method) **(Supplementary Fig. 5e,f)**. Crucially, MUC5AC and MUC5B are not functionally interchangeable—in murine models, MUC5B, but not MUC5AC, is essential for maintaining immune homeostasis and controlling infections of the upper airway, primarily through promoting mucocilliary clearance^55^—suggesting imbalances among these cell types could have profound effects on host defense.

We next sought to leverage the anatomical bifurcation present in our cohort to understand the functional differences associated with less (non-polyp) and more (polyp) severe T2I inflammatory disease **(Fig. 2d)**. Having observed striking differences within specific cell clusters driven by genes strongly associated with polyposis **(Fig. 2b,c)**, we quantified the numerical over-representation of cells from non-polyp and polyp ecosystems within each cluster and type **(Supplementary Table 3)**. The clusters composing basal, differentiating/secretory, and glandular cell types showed the most significant links to disease-state (p-values by Fisher’s with least significant difference**; Fig. 2d; Supplementary Table 3)**. Nasal polyps have been hypothesized to arise from the presence of opportunistic or pathogenic microorganisms^8^; thus, we first compared the transcriptomes of differentiating/secretory cells (containing *KRT8* secretory and goblet cells) **(Fig. 2e; Supplementary Table 3)**, which typically serve important antimicrobial functions through secreted products^4^. In non-polyp epithelium, we detected robust expression of antimicrobials (*MSMB, SCGB1A1, STEAP4, PSCA*, and *LYPD2*) which were diminished in polyp epithelium **(Fig. 2e)**. In turn, secretory cells recovered from polyps expressed *CST4* and *CST1,* associated with inhibition of protease activity^56^, and *IGFBP3, TFF3*, and *EGLN3,* indicating attempts at curtailing tissue growth^57^ and promoting tissue tolerance and repair^5,58,59^ **(Fig. 2e)**. Thus, secretory cells from polyps appear to supplant antimicrobial functions with tissue-reparative ones. No comparison between polyp and non-polyp glandular and ciliated cells was possible, as these clusters were nearly absent from polyp ecosystems **(Fig. 2d; Supplementary Fig. 6a; Supplementary Table 3)**.

As basal progenitor cells give rise to the specialized cell types present in the epithelium^3,10^, and glandular and ciliated cells were significantly reduced among polyps **(Fig. 3a, Supplementary Fig. 6a)**, we sought to more formally examine the distribution of epithelial cell types/states present in each sample. Appropriate biodiversity is key to the robustness of any ecological system^60^. Thus, we used Simpson’s index of diversity^61^—a measure of the richness of an ecosystem—to characterize the composition of epithelial cells across basal, differentiating/secretory, glandular, and ciliated groupings in the non-polyp and polyp ecosystems, as this metric accounts for both the number of distinct cell types present (e.g. species) and the evenness of composition across those cell types (e.g. relative abundance of species to each other**; Methods)**. Our data indicate a significant loss of epithelial diversity in nasal polyps, largely stemming from the aforementioned decrease in glandular and ciliated cells, and an enrichment in basal cells **(Fig. 3b),** that tracks the spectrum observed via clinical diagnoses **(Supplementary Fig. 6b)**. Furthermore, the rank-ordered pathological assessment of our patient tissue samples positively correlated with basal cell frequency (r=0.6252, p<0.0022) and negatively with epithelial diversity (r=-0.6824, p<0.0009; **Supplementary Fig 6c**). Intriguingly, when calculating Simpson’s index of diversity across parenchymal cells alone (e.g. fibroblast, endothelium, immune), or the entire cellular ecosystem for each patient, we observed a reduced index of diversity in polyps for parenchymal cells, but an overall increase across all cell types **(Supplementary Fig. 6d)**. Given the relative distribution of cell types present **(Supplementary Fig. 6a)**, and how evenness factors into this equation, we speculate that the immune cells in polyps may represent an overcorrection in attempting to restore balance to the epithelial compartment.

**Figure 3.**
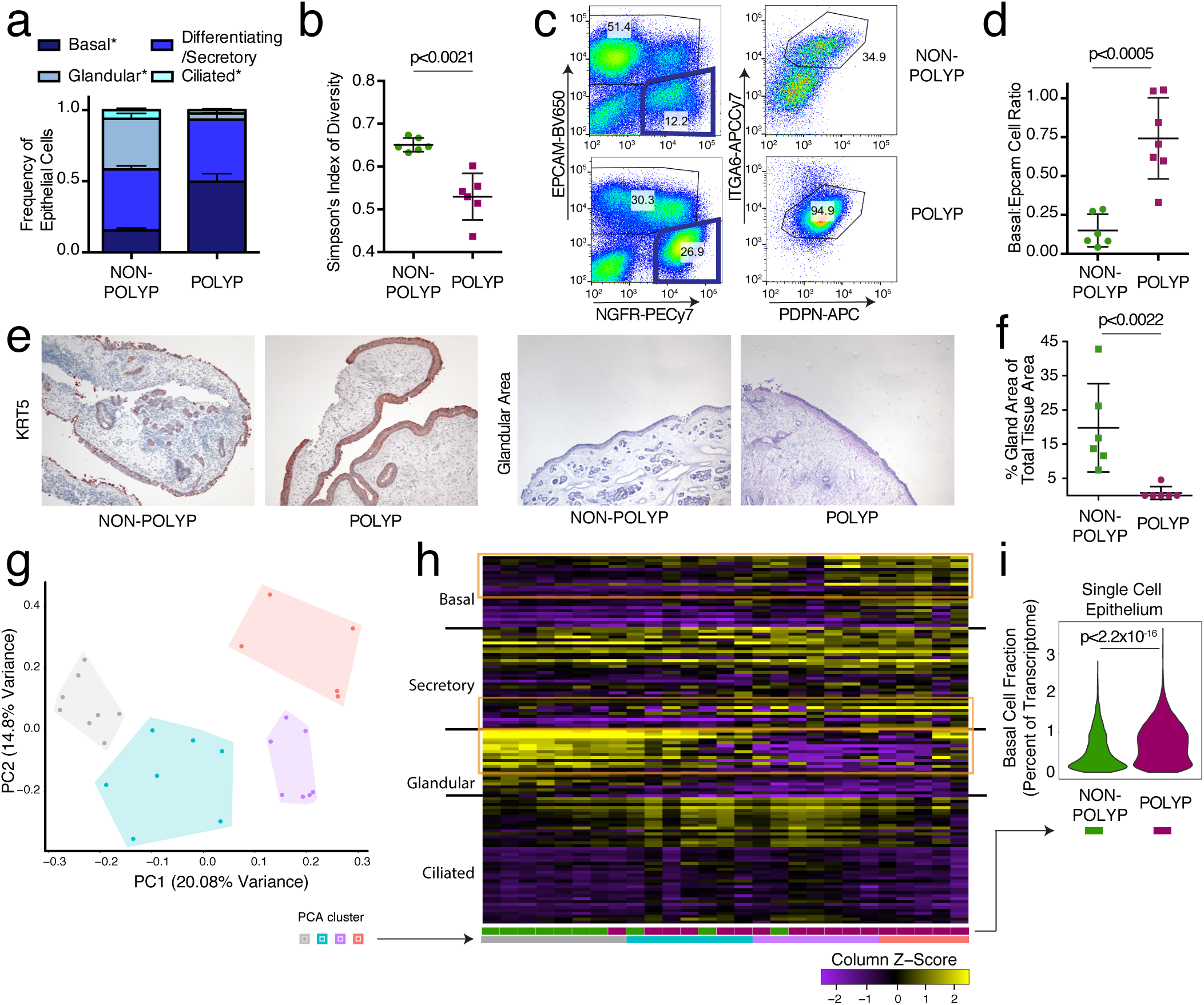
The ecological diversity of airway epithelial cells is reduced and basal cells are significantly increased in individuals with nasal polyps. **(a)** The frequency of each cell subset amongst all epithelial cells recovered in scRNA-seq calculated for each sample; n= 6 non-polyp, 6 polyp samples, *t-test p<0.05 for indicated cell types, non-polyp vs polyp. **(b)** Simpson’s index of diversity over epithelial cell types, an indication of the total richness present within an ecosystem, calculated for each sample; points represent individual samples, *t-test p<0.0021. **(c)** Representative gating strategy of flow cytometry performed on single cell suspensions of cells used to identify apical epithelial cells and basal cells (see **Supplementary Figure 7** for full gating strategy). **(d)** Quantification of flow cytometry for the ratio of basal to Epcam+ epithelial cells recovered from tissue; points represent individual samples; n = 6 non-polyp, 7 polyp samples, *t-test, p<0.0005. **(e)** Representative histology for KRT5 (basal cell marker) and for glandular area quantification on sections from non-polyp or polyp patients; 5x magnification. **(f)** Quantification of the glandular area detected in haematoxylin and eosin stained tissue sections from non-polyp or polyp patients; points represent individual patients; n = 6 non-polyp, 6 polyp samples, *t-test, p<0.0022. **(g)** Principal components analysis (PCA) over epithelial subset-specific genes (**Supplementary Table 3**) on 27 whole tissue RNA-seq samples from non-polyp (n=9), or polyp (n=18) patients with clusters identified by KNN (n= set to 4; **Methods**). **(h)** Column-normalized heatmap over epithelial subset-specific genes (Supplementary Table 3) grouped by KNN-clusters from **Fig. 3g**, color bars disease annotation (non-polyp= green, polyp= purple) and lower bar clusters from **Fig. 3g**. **(i)** Violin plot of expression contribution to a cell’s transcriptome of basal cell genes (effect size 0.457, polyp vs non-polyp; see **Methods** and **Supplementary Table 4**) across all recovered epithelial cells in non-polyp and polyp ecosystems; n= 6 non-polyp, 6 polyp samples, *t-test p<2.2x10^−16^.

To confirm our epithelial findings, we utilized several approaches. First, leveraging a flow cytometric gating strategy for human basal cells from the published literature^10^ **(Fig. 3c; Supplementary Fig. 7)**, we found that the frequency of basal cells is significantly increased in polyps at the expense of differentiated epithelial cells in an orthogonal cohort of 13 additional patients **(Fig. 3d)**. Second, using histology to ensure that our scRNA-Seq and flow results could not be explained as a dissociation induced artifact, we confirmed a thickening of the KRT5-positive region and a striking loss of glands in polyps, in direct accordance with our scRNA-seq data **(Fig. 3e,f; Methods)**. Finally, utilizing marker genes for specialized lineages **(Supplementary Table 3),** we deconvolved bulk tissue RNA-seq of another orthogonal cohort (n=27 patients) to look more globally at intact tissues (which, without curation, were largely driven by immune signatures**; Supplementary Fig. 2)**. Focusing on genes that define human basal, secretory, glandular, and ciliated cell subsets, we identified four clusters of patients (K-nearest neighbors (KNN) on a principal components analysis (PCA)**; Fig. 3g,h; Methods)**: a non-polyp cluster (grey) enriched in secretory and glandular signatures, and then three increasingly polyp enriched ones that showed more pronounced basal and ciliated cell programs (cyan), then lost glandular ones (lilac), and eventually, in the most severe cases, lost core ciliated genes (coral, as determined by number of previous surgeries, time to polyp regrowth, and eosinophilia) **(Fig. 3h; Supplementary Tables 1 & 3; Methods)**. Within polyps, we also confirmed upregulation of the tissue reparative program observed in **Fig. 2e** for differentiating/secretory cells. Serial profiling starting at incipient disease will be required to investigate whether CRS is divisible into a strict clinical divergence, or if individuals with polyps traverse a disease landscape from healthy tissue through that observed in non-polyps en route to polyposis.

To address if basal cell hyperplasia and a loss of secretory cell function characterizes deviations from healthy tissue, as well as from the CRS non-polyp disease state, we used two publically-available RNA-seq datasets containing normal human sinus mucosal biopsies, and non-eosinophilic and eosinophilic nasal polyps^8,62,63^ **(Methods).** Re-analyzing each sample for the fraction of basal or secretory cell markers present amongst genes representative of those two lineages, we identify a significantly increased basal and decreased secretory cell fraction in polyp tissues relative to healthy controls **(Supplementary Fig. 7c,d; Supplementary Table 3)**. This mirrors the findings from our cohort between non-polyp and polyp tissue, highlighting that the changes in cellular composition we observe also typify the divergence between the healthy and polyp states. Taken together, these data suggest that nasal polyps, relative to both healthy and CRS non-polyp tissue, are comprised of a barrier with diminished function amongst secretory cells, an almost complete loss of glandular cells, and an enhanced basal cell state throughout **(Fig. 3i; Supplementary Table 4)**.

We next sought to understand what mechanisms might account for the decreased epithelial diversity in polyps. Basal cells are the stem cells for upper airway respiratory epithelium^3,10^, though evidence suggests that in severe injury, trans- and de-differentiation may also account for tissue repair^52^. By comparing the transcriptomes of basal cells between our polyp and non-polyp ecosystems **(Fig. 4a)**, we identified elevated polyp expression of a set of transcripts— including *POSTN, PTHLH, ALOX15, SERPINB2, HS3ST1, CDH26, MMP10* and *CCL26*— involved in extracellular matrix remodeling and chemo-attraction of effector cells^46,64^, along with a decrease in protease inhibitor expression (*SPINK5*)^65^ and metabolic genes (*ALDH3A1, CLCA4, GLUL*) **(Fig. 4a).** As several of these genes are known to be IL-4/IL-13 responsive^66^, we looked more globally at gene sets induced by these cytokines across basal and other epithelial subsets. We determined that a combined IL-4/IL-13 signature **(Fig. 4b,c; Supplementary Fig. 8a; Supplementary Table 4)** is strongly induced not only in differentiated polyp epithelium, but also at the level of basal cells, and has a striking effect size between disease states **(Fig. 4c)**. Conversely, Type-I IFN and IFNγ signatures—indicative of a Type-1 immune module^14,66^—have small effect sizes **(Supplementary Fig. 8b; Methods, Supplementary Table 4)**. Furthermore, from specific hallmark genes, we observed that the balance of WNT (*CD44*) and Notch (*HEY1*) signaling appears altered in epithelium from polyps in favor of WNT **(Fig. 4a)**. We further confirmed this across full gene sets for these pathways^50,51,67^ **(Fig. 4c; Supplementary Table 4)**. Such aberrant signaling may contribute to failed basal cell differentiation.

**Figure 4.**
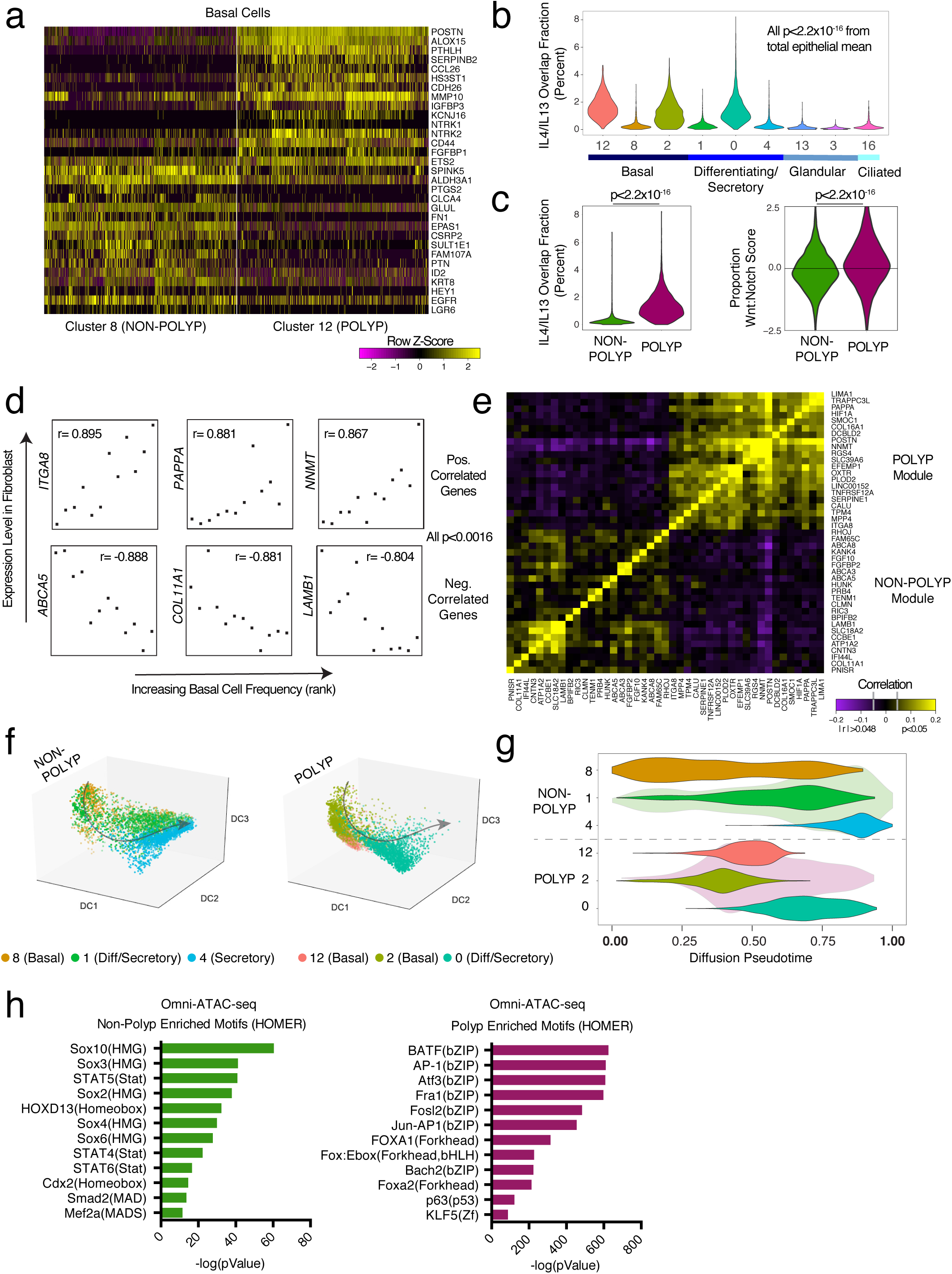
Extrinsic pathways converge at the chromatin level in basal cells to intrinsically impair differentiation. **(a)** Row-normalized heatmap of select differentially expressed genes using bimodal test over single-cells from basal cell clusters 8 and 12; n= 6 non-polyp, 6 polyp samples (see Supplementary Table 3 for genes and statistics, *bimodal test, all displayed genes p<1.97x10^−39^ or less with Bonferroni correction for multiple hypothesis testing based on number of genes tested). **(b)** Violin plot of expression contribution to a cell’s transcriptome over IL-4/IL-13 commonly-induced gene signature in respiratory epithelial cells; n= 6 non-polyp, 6 polyp samples (see **Methods**, **Supplementary Figure 8** for unique genes, and **Supplementary Table 4** for gene lists used, all p<1.98x10^−15^, relative to mean score, with Bonferroni correction for multiple comparisons). **(c)** Violin plots of shared IL-4/IL-13 signature expression contribution to a cell’s transcriptome (effect size 1.305, polyp vs. non-polyp) and of the Wnt:Notch target gene proportion (effect size 0.334, polyp vs. non-polyp, NB: axis truncated at −2.5 and 2.5) (see **Methods**, and **Supplementary Table 4** for gene lists used, zero indicates equal scores, Wnt-pos, Notchneg direction) across all epithelial cells grouped by disease state; n= 6 non-polyp, 6 polyp samples, *t-test p<2.2x10^−16^ for both. **(d)** Selected genes detected in fibroblasts from single-cell data which correlate with the samples ranked by basal cell frequency detected within each ecosystem; n= 6 non-polyp, 6 polyp samples, all genes used: abs(r)> 0.7651, p<0.0037. **(e)** A clustered correlation matrix of genes identified in d) in single-cell data from fibroblasts; abs(r)> 0.048 is p<0.05 significant based on asymptotic p-values. **(f)** Pseudotime analysis using diffusion mapping (see **Methods**) of selected clusters of epithelial cells, colored by cluster; n= 3,516 cells (clusters 8/1/4), n= 4,064 cells (clusters 12/2/0), and n= 6 non-polyp, 6 polyp samples, diffusion map and DC (diffusion coefficients) are calculated over the set of basal and apical marker genes identified in **Fig. 1a**, see **Supplementary Table 3.** **(g)** Violin plot of pseudotime component (see **Methods**) for cells in respective clusters; shading in green is non-polyp distribution and in purple is polyp distribution underlying respective clusters. **(h)** Omni-ATAC-seq (see Methods) profiling and HOMER motif enrichment of non-polyp peaks over all peaks as background and polyp peaks over all peaks as background on low-input sorted basal cell populations; n= 3 non-polyp and n=7 polyp; all q-value < 0.0002 Benjamini corrected.

We next sought to contextualize our basal cell findings related to extracellular matrix remodeling within their larger cellular ecosystems by asking whether cells which compose the basal cell niche, such as fibroblasts^68^, were altered in polyps potentially contributing to basal cell dysfunction. Thus, we looked for genes in fibroblasts whose expression correlated (all genes: abs(r)>0.7651, p<0.0037) with basal cell frequency **(Fig. 4d; Supplementary Fig. 6b; Methods)**. Clustering over fibroblast genes that either positively or negatively correlated with basal cell frequency in our single-cell data identified a polyp-enriched gene module in fibroblasts associated with *ITGA8*, a hallmark gene of these cells during development of new airways but also fibrosis **(Fig. 4e)**^69,70^. Taken together, our data suggest that basal cells harbor an imprint of IL-4/IL-13 signaling, and are found amongst an altered niche constituent.

While we identified a fundamental loss of glandular and ciliated cells in polyps, we recovered equivalent (but functionally altered) proportions of differentiating/secretory cells. Thus, we asked if by reconstructing how basal cells transition to mature secretory cells within non-polyp or polyp tissue, we could gain insight into the mechanisms leading to the observed loss in cellular diversity and function. Using diffusion pseudotime^71^ mapping, which seeks to provide the most likely reconstruction for the developmental progression of a set of cells **(Methods)**, we built a trajectory for cells within basal and differentiating/secretory epithelial clusters (non-polyp clusters: 8-basal, 1-differentiating/secretory, 4-secretory; and polyp clusters: 12-basal, 2-basal, 0-differentiating/secretory; running several iterations starting from a random seed cell in cluster 8), over the combined basal and apical marker gene list **(Fig. 4f; Supplementary Fig. 9a, Supplementary Table 3).** By calculating a pseudotime trajectory for cells from both non-polyps and polyps together, we were then able to ask where cells from each disease state fall along an inferred temporal axis **(Fig. 4f,g, Supplementary Fig. 9a)**. In the non-polyp ecosystem, we observed that basal cells (cluster 8) traverse a much wider swath of pseudotime, with the majority of cells distributed towards the end of the trajectory (cluster 4**; Fig. 4f,g; Supplementary Fig. 9a)**. Conversely, in polyps, basal cell clusters 2 and 12 accumulate shy of the midpoint of the trajectory, failing to productively contribute to later differentiation states **(Fig 4f,g; Supplementary Fig. 9a).** Furthermore, cluster 0 does not reach the terminally differentiated state observed in cluster 4, suggesting that the aforementioned expression differences **(Fig. 2d,e)** may, in part, be due to a failure of terminal differentiation. Ordering our cells according to this common axis, we examined our full gene expression matrix to identify genes that exhibited a significant change in correlation between non-polyps vs. polyps and thus might become dysregulated in polyps during differentiation. We found that: (1) DLK2, DLL1, JAG2, DKK3 (mediators of Wnt and Notch signaling, **Fig. 4c**) and (2) POSTN, FN1, TNC (extracellular matrix components, FN1 and TNC are ligands for ITGA8; **Fig. 4e**) were both significantly negatively correlated with pseudotime in non-polyps, and had altered correlations in polyps (based on abs(Fisher’s Z) > 3.8**; Supplementary Fig. 9b; Supplementary Table 3** for full list and statistics). Future work will seek to rigorously test how these deviations in pathways observed in polyps affect differentiation trajectories in controlled non-human systems.

As these data suggested an intrinsic impairment in the differentiation of basal cells in polyp tissue, we next sorted basal cells **(Supplementary Fig. 7)** from 4 non-polyp and 7 polyp tissues (3 non-polyp and 7 polyp retained through data quality filtering) and performed Omni-ATAC-Seq to profile chromatin accessibility^72,73^ **(Methods)**. Chromatin exists in a poised state in stem cells^74^, providing a form of epigenetic memory^75^, with influence on the propensity to differentiate towards particular cell fates. Among our data, we identified that polyp basal cells had an enrichment in peaks (**Methods**) for bZIP transcription factor target motifs, including various AP-1 family members, JUN, FOXA1, BACH2, and p63 itself (*TP63* and *JUNB* were significantly negatively correlated with a productive differentiation trajectory in non-polyps, with a deviation observed in polyps**; Supplementary Fig. 9b**), **(Fig. 4h; Supplementary Table 5)**. These proteins have previously been associated with the maintenance of an undifferentiated state, chromatin opening, and oncogenesis^76^. Conversely, accessible motifs in non-polyp basal cells were enriched for Sox, STAT, and Mef2 family members, typically implicated in productive development, differentiation and regeneration of tissues^77^ **(Fig. 4h; Supplementary Table 5)**. Clustering of enriched motifs in non-polyp samples revealed Sox2/Sox4/Sox6 and Sox3/Sox10 modules **(Supplementary Fig. 10a,b)**; both decreased in polyps **(Supplementary Fig. 10b)**. Furthermore, in polyp basal cells, p63 shows reduced correlation with the Sox and AP-1 modules, suggesting that higher-order changes in chromatin structure may affect the activity of this key basal cell transcription factor in severe disease^78^ **(Supplementary Fig. 10)**. Together, our differentiation trajectory analysis and epigenetic studies suggest that during chronic T2I, basal cell differentiation is intrinsically impaired through the influence of extrinsic cues (e.g., IL4/IL13 and niche components). This results in a barrier with reduced cellular and functional diversity, akin to an ecosystem with compromised biodiversity, and consequently an overall reduction in tissue health^60^.

One fundamental goal of understanding the cellular and molecular pathways activated in T2I is to provide mechanisms that may explain the dramatically increased incidence of allergic inflammatory disease, particularly beyond conventional immune mechanisms^79^. While a signature of immune cell infiltration dominates bulk transcriptomic analyses of polyps relative to non-polyp tissue in CRS **(Supplementary Fig. 2),** by utilizing scRNA-seq in combination with a well-defined patient cohort across the severity spectrum of a human chronic allergic inflammatory disease, we have uncovered previously obscured biological insights into the concerted cellular shifts that occur in the epithelial compartment, and their functional consequences. Our data may help to explain why nasal polyposis is associated with recurrent infections by specific microorganisms^8^, and how a monoclonal antibody targeting the shared IL-4/IL-13 receptor can reduce nasal polyp burden^80^. We show striking differences in the antimicrobial peptides produced by secretory cells, a loss of glandular cells producing *MUC5B*^55^, and that IL-4/IL-13 strongly induce a transcriptional program already at the level of basal progenitor cells^81^. Our results suggest that therapeutic modulation of basal cell differentiation during chronic inflammation could help to restore the homeostatic balance. Taken together with recent studies of the murine intestinal tract and skin^17,82-84^, our work provides human evidence for the emerging paradigm of stem cell dysfunction altering the set point of a barrier tissue. In future work, it will be of interest to determine the relative contributions of memory stored in distinct compartments: by cells classically viewed to drive allergic inflammation, such as T and B cells, and within the epithelium itself^85^. While the immune response in T2I may represent an attempt to re-set appropriate biodiversity within a tissue **(Fig. 3b, Supplementary Fig. 6d)**, based on our results, we propose that basal cells can form “memories” of chronic exposure to an inflammatory T2I environment, shifting the entire specialized cellular ecosystem away from productive differentiation, and propagating disease.

## METHODS

### Study Participants and Design for Single-Cell Study

Subjects between the ages of 18 and 75 years were recruited from the Brigham and Women’s Hospital (Boston, Massachusetts) Allergy and Immunology clinic and Otolaryngology clinic between May 2014 and August 2017 **(Supplementary Table 1)**. The Institutional Review Board approved the study, and all subjects provided written consent. Tissue was collected at the time of elective endoscopic sinus surgery from patients with physician-diagnosed CRS with and without nasal polyps based on established guidelines^86^. Patients with polyps include both aspirin-tolerant chronic rhinosinusitis with nasal polyps (CRSwNP) and individuals with aspirin-exacerbated respiratory disease (AERD). Patients were suspected of having AERD if they had asthma, nasal polyposis, and a history of respiratory reaction on ingestion of a COX 1 inhibitor, with confirmation via a graded oral challenge to aspirin. Subjects with cystic fibrosis and unilateral polyps were excluded from the study. No distinctions were made between these two groupings in our study as both present with polyposis, but we present the information of clinical diagnosis in **Supplementary Table 1**.

Tissue segments (one per patient) for bulk tissue RNA-seq was immediately placed in RNAlater (Qiagen) for RNA extraction, and for patient samples loaded on Seq-Well and for flow-sorting to ATAC-seq, tissue was received in-hand, placed in RPMI (Corning) with 10% FBS (ThermoFisher 10082-147) and immediately on ice for transport. Details of the subjects’ characteristics included in scRNA-seq cohort, tissue RNA-seq cohort, and flow cytometry/ATAC-seq cohort (including age, gender, medication use, and disease severity) are included in **Supplementary Table 1**.

NB: Originally, we enrolled two healthy control subjects with no history of CRS or nasal polyposis who were undergoing sinus surgery for concha bullosa. However, these two subjects upon pathology evaluation were noted to have mild eosinophilia, a chart review revealed a history of allergic rhinitis and asthma, and their diagnosis was updated to CRSsNP clinically by the surgeon upon follow-up visits so we updated their status accordingly in our study. Additionally, non-polyp patient 6 was sampled twice (denoted as 6A and 6B), representing distinct cells that were captured on two different Seq-Well arrays. As such, they should not be viewed as a technical replicate.

### Tissue Digestion

Single-cell suspensions from collected surgical specimens were obtained using a modified version of a previously published protocol^87^, described below in detail. Each specimen was received directly in hand and processed directly with an average time from patient to loading onto the SeqWell platform of 3 total hours, and never exceeding 4 hours. Surgical specimens were collected into 30 mL of ice cold RPMI (Corning). Specimens were finely minced between two scalpel blades and incubated for 15 minutes at 37°C in a rotisserie rack with end-over-end rotation in 25 mL digestion buffer supplemented with 600 U/mL collagenase IV (Worthington) and 20 ug/mL DNAse 1 (Roche) in RPMI with 10% fetal bovine serum. After 15 minutes, samples were triturated five times using a syringe with a 16G needle and returned to the rotisserie rack for another 15 minutes. At the conclusion of the second digest period, samples were triturated an additional five times using a syringe with a 16G needle, at which point the digest process was stopped via the addition of EDTA to 20mM. Samples were typically fully dissociated at this step and were filtered through a 70uM cell strainer and spun down at 500G for 10 minutes followed a rinse with ice-cold PBS (ThermoFisher 10010023, Ca/Mg free) to 30 mL total volume. RBCs were lysed using ACK buffer (ThermoFisher A1049201) for 3 minutes on ice to remove red blood cells, even if no RBC contamination was visibly seen in order to maintain consistency across patient groups. Cells were then washed with sterile PBS and spun down at 500G for 5 minutes, resuspended in complete RPMI medium with 2% FCS (RPMI1640 [ThermoFisher 61870-127], 100 U/ml penicillin [ThermoFisher 15140-122], 100 ug/mL streptomycin [ThermoFisher 15140-122], 10 mM HEPES [ThermoFisher 15630-080], 2% FCS [ThermoFisher 10082-147], 50 ug/mL gentamicin [ThermoFisher 15750-060]), and counted to adjust concentration to 100,000 cells/mL for loading onto SeqWell arrays.

### Flow Cytometry, Cell Sorting, and Analysis

Single-cell suspensions in FACS Buffer (HBSS [ThermoFisher 14170161, Ca/MgFree] supplemented with 2% FCS) were pre-incubated with Fc Block before staining for surface antigens. The following antibodies were used to identify basal cells via flow cytometry: FITC anti-human THY1 (Biolegend, clone 5E10), Brilliant Violet 421 anti-human CD45 (Biolegend, clone HI30), Brilliant Violet 650 anti-human EPCAM (Biolegend, clone 9C4), APC/Cy7 anti-human ITGA6 (Biolegend, clone GoH3), PE/Cy7 anti-human NGFR (Biolegend, clone ME20.4), APC anti-human PDPN (Biolegend, clone NC-08). Cells were stained for 30 minutes on ice in FACS buffer and then washed for immediate sorting. Cells were sorted on a BD FACSAria Fusion cell sorter using BD FACSDiva software. Up to 10,000 Basal cells were sorted into 100 µL BAM banker (Wako chemicals) for ATAC sequencing and cooled to −80 C using a “Mr. Frosty” freezing container (Thermo scientific). Samples were stored at −80 until thawed for omni-ATAC-seq. FlowJo v10 by TreeStar was used to generate plots.

### Histologic analyses

Biopsies were fixed in 4% paraformaldehyde, embedded in paraffin, and 6 µm sections were prepared and stained with hematoxylin and eosin for quantification of glandular areas. Photomicrographs encompassing the entire area of each biopsy were taken. Total and glandular areas were measured with Image J software and expressed as glandular area as a percentage of total area. Sections for immunohistochemical staining were incubated with 3% hydrogen peroxide to inactive endogenous peroxidases, Dako Target Retrieval Solution pH 9, and rabbit anti-human KRT5 polyclonal Ab (GeneTex) or non-immune rabbit IgG. Antibody binding was visualized with the DAKO EnVision+ System-HRP (AEC) system, and sections were counterstained with Gill’s hematoxylin number 2 (Sigma). Sections incubated with non-immune rabbit IgG were negative.

### Single-cell RNA-sequencing

Once a single-cell suspension was obtained from freshly resected sinus tissue, we utilized the Seq-Well platform for massively parallel scRNA-seq to capture transcriptomes of single cells on barcoded mRNA capture beads. Full methods on implementation of this platform are available in Gierahn et al.^11^. Briefly, 20,000 cells were loaded onto one array preloaded with barcoded mRNA capture beads (ChemGenes). The loaded arrays containing cells and beads were then sealed using a polycarbonate membrane with a pore size of 0.01 µm, which allows for exchange of buffers but retains biological molecules confined within each nanowell. Subsequent exchange of buffers allows for cell lysis, transcript hybridization, and bead recovery before performing reverse transcription en masse. Following reverse transcription using Maxima H Minus Reverse Transcriptase (ThermoFisher EP0753) and an Exonuclease I treatment (NewEngland Biolabs M0293L) to remove excess primers, PCR amplification was carried out using KAPA HiFi PCR Mastermix (Kapa Biosystems KK2602) with 2,000 beads per 50 uL reaction volume. Libraries were then pooled in sets of six (totaling 12,000 beads) and purified using Agencourt AMPure XP beads (Beckman Coulter, A63881) by a 0.6X SPRI followed by a 0.7X SPRI and quantified using Qubit hsDNA Assay (Thermo Fisher Q32854). Quality of WTA product was assessed using the Agilent hsD5000 Screen Tape System (Agilent Genomics) with an expected peak >1000bp tailing off to beyond 5000bp, and a small/non-existent primer peak, indicating a successful preparation. Libraries were constructed using the Nextera XT DNA tagmentation method (Illumina FC-131-1096) on a total of 600 pg of pooled cDNA library from 12,000 recovered beads using index primers with format as in Gierahn et al^11^. Tagmented and amplified sequences were purified at a 0.6X SPRI ratio yielding library sizes with an average distribution of 650-750 base pairs in length as determined using the Agilent hsD1000 Screen Tape System (Agilent Genomics). Two arrays were sequenced per sequencing run with an Illumina 75 Cycle NextSeq500/550v2 kit (Illumina FC-404-2005) at a final concentration of 2.8pM. The read structure was paired end with Read 1 starting from a custom read 1 primer^11^ containing 20 bases with a 12bp cell barcode and 8bp unique molecular identifier (UMI) and Read 2 containing 50 bases of transcript information.

### Single-cell RNA-sequencing computational pipelines and analysis

Read alignment was performed as in Macosko et al^88^. Briefly, for each NextSeq sequencing run, raw sequencing data was converted to demultiplexed FASTQ files using bcl2fastq2 based on Nextera N700 indices corresponding to individual samples/arrays. Reads were then aligned to hg19 genome using the Galaxy portal maintained by the Broad Institute for Drop-Seq alignment using standard settings. Individual reads were tagged according to the 12-bp barcode sequenced and the 8-bp UMI contained in Read 1 of each fragment. Following alignment, reads were binned onto 12-bp cell barcodes and collapsed by their 8-bp UMI. Digital gene expression matrices (e.g. cells-by-genes tables) for each sample were obtained from quality filtered and mapped reads, with an automatically determined threshold for cell count. UMI-collapsed data was utilized as input into Seurat^22^ (https://github.com/satijalab/seurat) for further analysis. Before incorporating a sample into our merged dataset, we individually inspected the cells-by-genes matrix of each as a Seurat object.

For analysis of all sequenced samples, we merged UMI matrices across all genes detected in any condition and generated a matrix retaining all cells with at least 500 UMI detected (n= 19,196 cells and 31,032 genes). This table was then utilized to setup the Seurat object in which any cell with at least 300 unique genes was retained and any gene expressed in at least 5 cells was retained (**Supplementary Information**: an R Script is included from this point to set up Seurat object and walk reader through dimensionality reduction and basic data visualization). The object was initiated with log-normalization, from a UMI+1 count matrix, scaling, and centering set to True. The total number of cells passing these filters captured across all patients was 18,624 cells with 22,575 genes, averaging 1,503 cells per sample with a range between 789 cells and 3,109 cells. Before performing dimensionality reduction, data was subset to include cells with less than 12,000 UMI, and a list of 1,627 most variable genes was generated by including genes with an average normalized and scaled expression value greater than 0.13 and with a dispersion (variance/mean) greater than 0.28. We then performed principal component analysis over the list of variable genes. For both clustering and t-stochastic neighbor embedding (tSNE), we utilized the first 12 principal components, as upon visual inspection of genes contained within, each contributed to a non-redundant cell type and this reflected the inflection point of the elbow plot. We used FindClusters (which utilizes a shared nearest neighbor (SNN) modularity optimization based clustering algorithm) with a resolution of 1.2 and tSNE set to Fast with the Barnes-hut implementation to identify 21 clusters across the 12 input samples.

### Cell Type Identification and within Cell Type Analysis

To identify genes which defined each cluster, we performed a ROC test implemented in Seurat with a threshold set to an area under the curve of 0.65. Top marker genes with high specificity were used to classify cell subsets into cell types **(Fig. 1a,b,c)** based on existing biological knowledge. Three clusters were considered doublets (n=588 cells) based on co-expression of markers indicative of distinct cell types at ~1/2 the expression level detected in the parent cell cluster (e.g. T cell and myeloid cell) and removed from further analyses yielding a matrix with 18,036 cells used in all subsequent steps. Closely related clusters were merged to cell types based on biological curation and analysis of hierarchical cluster trees yielding ten total cell types **(Fig. 1a,b,c)**. We identified a much smaller number of eosinophils than expected in our single-cell data. However, our own experience in attempting to isolate RNA from eosinophils or eosinophil-rich tissue (such as eosinophilic esophagitis) indicates that their RNA is quite labile. Specifically, if we do not place tissue immediately into RNA-later within 10 minutes, we cannot reliably detect eosinophil associated transcripts, and preliminary SeqWell experiments on patients with active eosinophilic esophagitis also yielded low capture of eosinophils (data not shown). However, flow cytometrically we recover from 0.5% to 5% of total cells fitting eosinophil profiles from polyps, and focused single-cell studies on granulocytes at the expense of the full ecosystem are possible and the topic of future work (data not shown). We also did not find a distinct cluster of ILCs as they are around 0.01 to 0.1% of CD45 cells across the CRS spectrum per existing literature^24^ and extrapolating to the number of CD45 cells we captured, we would have detected between 0.8 and 8 ILCs. To investigate further granularity present within cell types, such as T cells, we subset these cells from the Seurat object and re-ran dimensionality reduction and clustering **(Supplementary Fig. 3)**. The process used for clustering and subset identification was adapted for each cell type to optimize the parameters of variable genes, principal components, and resolution of clusters.

### Differential Expression and Fractional Contribution of Gene Set to Transcriptome

To identify differentially expressed genes within cell types across non-polyp and polyp disease states, we utilized the ‘bimod’ setting in FindMarkers implemented in Seurat based on a likelihood ratio test designed for single-cell differential expression incorporating both a discrete and continuous component^89^. To determine the expression contribution to a cell’s transcriptome of a particular gene list, we summed the total log-normalized expression values for genes within a “list of interest” and divided by the total amount of log-normalized transcripts detected in that cell, giving the proportion of a cell’s transcriptome dedicated to producing those genes. For comparison of Wnt and Notch signaling, we z-scored the expression contribution metric and subtracted the value of Notch from Wnt yielding a metric centered on zero if both scores are equivalent, or weighted in the positive direction if enriched in Wnt. For reference gene lists used, please see **Supplementary Table 4**.

### Simpson’s Index of Diversity, and Fibroblast Gene correlation with Basal Cell Frequency

To measure the “richness” of the epithelial ecosystem, we employed Simpson’s Index of Diversity (D), which we present as (1-D), and ranges between 0 and 1, with greater values indicating larger sample diversity^61^. This measure takes into account the total number of members of a cell type, the number of cell types, and the total number of cells present. We calculate (1-D) for each sample. To determine genes correlated in specific cell types (e.g. fibroblasts) with the frequency of basal cells present in a cellular ecosystem, we correlated the average log-normalized single-cell count data for each gene to the rank of samples determined by increasing frequency of basal cells in each ecosystem (8.2% to 19.1% for non-polyp and 27.9% to 70.1% for polyp samples, **Supplementary Fig. 6b**).

### Tissue RNA-sequencing

Population RNA-seq was performed using a derivative of the Smart-Seq2 protocol for single cells^90^. In brief, tissue was collected directly into RNAlater (Qiagen) in the surgical suite and stored at −80 until RNA isolation. RNA was isolated from 30 patients using phenol/chloroform extraction and normalized to 5ng as the input amount for a 2.2X SPRI ratio cleanup using Agencourt RNAClean XP beads (Beckman Coulter, A63987). After oligo-dT priming, Maxima H Minus Reverse Transcriptase (ThermoFisher EP0753) was utilized to synthesize cDNA with an elongation step at 52°C before PCR amplification (15 cycles) using KAPA HiFi PCR Mastermix (Kapa Biosystems KK2602). Sequencing libraries were prepared using the Nextera XT DNA tagmentation kit (Illumina FC-131-1096) with 250pg input for each sample. Libraries were pooled post-Nextera and cleaned using Agencourt AMPure SPRI beads with successive 0.7X and 0.8X ratio SPRIs and sequenced with an Illumina 75 Cycle NextSeq500/550v2 kit (Illumina FC-404-2005) with loading density at 2.2pM, with paired end 35 cycle read structure. Samples were sequenced at an average read depth of 7.98 million reads per sample and 3 samples not meeting quality thresholds were excluded from further analyses yielding 27 total useable samples.

### Tissue RNA-sequencing Data Analysis

Samples were aligned to the Hg19 genome and transcriptome using STAR^91^ and RSEM^92^. 3 samples were excluded for low transcriptome alignment (<25%), so we retained 27 samples for further analyses. Differential expression analysis was conducted using DESeq2 package for R^93^. Genes regarded as significantly differentially expressed were determined based on an adjusted p-value using a Benjamini-Hochberg adjusted p-value to correct for multiple comparisons with a false discovery rate <0.05. We performed Ingenuity Pathway Analysis (IPA, Qiagen) through an instance available through the Broad Institute on the top 1000 genes (all adjusted p<0.05) differentially expressed from our DESeq2 analysis, taking into account corresponding log-fold change for each gene. We also subset the tissue RNA-seq matrix based on genes found in **Supplementary Table 3**, which, from our single-cell marker discovery, were specific for basal, differentiating/secretory, glandular, or ciliated cells. We then ran PCA and KNN clustering implemented in R over these genes in order to identify the greatest vectors of variance across samples within the epithelial cell compartment.

For re-analysis of published data, we used two publically-available RNA-seq data sets: one profiling normal human olfactory mucosa and the other assessing differences in gene expression between healthy, non-eosinophilic nasal polyps and eosinophilic nasal polyps^8,62,63^ NB: analysis is done per sample and as such no comparisons across the data sets or samples are made.

### Diffusion Pseudotime Mapping for Differentiation Analysis

Diffusion pseudotime^71^ was calculated using the scanpy python package ‘dpt’ function on log normalized data separately for clusters 8, 1 and 4 (predominantly non-polyp, **Supplementary Table 3**) and 12, 2, and 0 (predominantly polyp, **Supplementary Table 3**) together by. A random root cell was chosen from cluster 8, as this was the basal cell cluster representative of the non-polyp (e.g. less aberrant) state, and we also ran iterations with random root cells chosen from the entire set of clusters and it assigned cluster 8 as the cluster most enriched at the beginning of the diffusion map, regardless. Plots were created with the seaborn, matplotlib, and pandas packages. Pearson correlations were then calculated for all genes in all cells tested, or for all genes in non-polyp cells and all genes in polyp cells, relative to pseudotime. Differential correlation testing was performed using the cocor package to identify significance for the difference between correlation coefficients using Fisher’s 1925 z-statistic, account for number of cells.

### Epigenetic Profiling of Basal Cells using Omni-ATAC-Seq

Accessible chromatin profiling using the Omni-ATAC-Seq protocol as described in Corces et al^73^ was performed on basal cells stored in 100 µL BAMBanker freezing media from 12 patients (n= 4 non-polyp and n= 8 polyp). Cells (ranging from 1,000 to 10,000) were thawed quickly in a 37°C rock bath and 900 µL of ice-cold PBS supplemented with Roche complete-Mini Protease inhibitor was added immediately. Cells were split into two 1.5mL Eppendorf DNA lo-bind tubes to serve as technical replicates. Cells were centrifuged at 500g for 5 minutes at 4°C, washed once in PBS with protease inhibitor, centrifuged at 500g for 5 minutes at 4°C and supernatant was removed completely using two separate pipetting steps with extreme caution taken to avoid resuspension (e.g. smooth and consistent aspiration). The transposition reaction consisted of 20µL total volume of the following mixture (10 µL 2X TD Buffer, 1 or 0.5 µL TDEnzyme, 0.1 µL of 2% digitonin, 0.2 µL of 10% Tween 20, 0.2 µL of 10% NP40, 6.6 µL of 1X PBS and 2.3 µL of nuclease free water). We performed replicates with two distinct concentrations of TDE since, when dealing with minute clinical samples, flow sorting can sometimes give variable cell numbers, and the ratio of TDE to cells is critical in determining the frequency with which cuts are made in the genome. We optimized in pilot experiments that for basal cell inputs in the range of 500 to 10,000 cells, the aforementioned two ratios gave expected patterns of nucleosome banding in gels (data not shown). We performed two reactions and then later, during in silico analysis, pooled peaks together for downstream analysis. The cells were resuspended into the transposition mixture and incubated at 37°C for 30 minutes in an Eppendorf Thermomixer with agitation at 300rpm. Transposed DNA was purified using a Qiagen MinElute Reaction Cleanup Kit with elution in 15 uL. Libraries were constructed from 10 µL of DNA using a 50 µL total reaction volume of NEB HF 2X PCR Master Mix with custom Nextera N700 and N500 index primers to barcode samples (also used in Smart-Seq2 protocol). We performed 14 cycles of PCR amplification and SPRI purified at 1.8X ratio. Based on the molarity of each library, we adjusted the number of subsequent PCR cycles to either 3, 4 or 5 more for each sample. We then performed a 0.25X reverse SPRI to remove larger fragments followed by a 1.7X SPRI to purify libraries for sequencing. Libraries were sequenced on an Illumina NextSeq with paired end 38 cycle read structure at a loading density of 1.95pM.

### ATAC-Seq Data Analysis

Reads were aligned using bowtie2 using the following flags: “-S -p 1 -X 2000 –chunkmbs 1000” then bams were created using samtools view with the following flags: ‘samtools view -bS -F 4 – “. Duplicates were removed with picard. Forward reads were shifted 4bp and negative reads were shifted 5bp using a custom python script and the pysam package as is recommended for ATAC-seq data. Samples for each patient were merged using samtools merge and all patients were downsampled to 3 million reads using custom python scripts and ‘samtools view’ with the ‘-b’ and ‘-s’ flags. MACS2 ‘callpeak’ command was used to call peaks on each sample with flags ‘“-f BAMPE’-q 0.001 –nomodel –shift −100 –extsize 200 -B –broad’. Peaks from all samples were merged into one peakfile with bedtools and counts of reads per peak for each sample was generated with bedtools multicov. DESeq2 was run with the design ~polyp, testing for significant differences between polyp and non-polyp samples on this peak matrix and differential peaks with Benjamini-Hochberg adjusted p-value less than 0.01 with ‘greater’ or ‘less’ null hypotheses were used in downstream analysis. Homer2 was run for known motif finding on differential peaks with the set of all peaks as background^94^. To determine a false discovery rate, Homer2 was run on sets of random peaks chosen with replacement from the set of all peaks.

### Statistical Analyses

Number of samples included in analyses are listed throughout figure legends and all represent distinct biological samples. The same surgeon performed surgeries on all individuals and was blinded to study design. No samples or cells meeting quality thresholds were excluded from analyses. Statistical analyses were performed using GraphPad Prism v7.0a, Seurat 1.4.0.1 implemented in RStudio, DESeq2 1.10.1 package implemented in RStudio, and Ingenuity Pathway Analysis run through the Broad Institute, and macs2, DESeq2 and Homer2 for omni-ATAC-Seq. All violin plots, which we elected to use due to zero-inflation in single-cell data, contain at minimum 498 individual data points in any one cluster (**Supplementary Table 3** for precise numbers of cells per cluster and type), and have points suppressed for ease of legibility. Violins are generated through default code implemented in Seurat for generation. Effect sizes for cell scores are displayed to complement t-tests. Unpaired 2-tail t-tests for direct comparisons and t-test with Holm-Sidak correction, Bonferroni correction, or Benjamini-Hochberg for multiple comparisons, depending on software package used, where appropriate. Pearson correlation thresholds were determined as significant through determination of asymptotic p-values through use of rcorr function in Hmisc, but exact corrected p-values by Holm-Sidak method for multiple comparisons are calculated for those highlighted in text using RcmdrMisc package. Comparison of Pearson correlation coefficients in pseudotime analyses was done using Fisher’s 1925 z-statistic account for the number of cells.

### Data Availability Statement

Marker gene lists for cell types identified in **Fig. 1a,b,** and from resultant analyses in **Fig. 2b,** and selected comparisons of differential expression in **Fig. 2e** and **Fig. 4a** are available as **Supplementary Information Tables** upon request. FASTQ file format data will be available shortly through dbGaP. The cells-by-genes matrix generated and analyzed during the current study will be available along with the published peer-reviewed manuscript as **Supplementary Information** along with an R code for standard implementation of Seurat.

## Supplementary Information

Is available upon request and consists of:

**Supplementary Table 1:** Patient information

**Supplementary Table 3:** Cell type specific marker genes, epithelial marker genes, and differential expression full gene lists presented in manuscript

**Supplementary Table 4:** Gene lists derived from literature and databases used for scoring

**Supplementary Table 5:** Omni-ATAC-Seq enriched motifs in basal cells

Will be made available upon peer-review and publication:

**Supplementary Table 2:** Cells-by-genes digital gene expression matrix

**R Script:** Loading cells-by-genes matrix into R and initiating Seurat object as illustrated in Fig. 1

## Acknowledgements

We thank S.L. Carroll and H. Raff for technical support in performing Seq-Well experiments and RNA extraction, J. Lai for histology support, L. Ludwig, J. Hammelman, and J. Buenrostro for reagents and analysis advice for ATAC-Seq, D. Lingwood, U.H. von Andrian, B. Walker, N. Yosef, S. Rakoff-Nahoum, S. Beyaz, C. Borges, M.B. Cole, N. Yosef, R. Satija, for discussions and comments on the manuscript, and members of the Shalek Lab for experimental and computational advice, and M. Morrison for administrative support. A.K.S. was supported by the Searle Scholars Program, the Beckman Young Investigator Program, NIH grants 1DP2OD020839, 2U19AI089992, 1U54CA217377, P01AI039671, 5U24AI118672, 2RM1HG006193, 1R33CA202820, and Bill and Melinda Gates Foundation grants OPP1139972 and BMGF OPP1116944; N.A.B. by NIH R01 HL120952 and Steven and Judy Kaye Young Innovators Award; T.M.L. by NIH R01 HL128241; J.A.B. by NIH U19 AI AI095219; D.F.D. by T32 AI007306 (to J.A.B.); K.M.B. by NIH AADCRC Opportunity Fund Award U19AI070535. J.O.M. is an HHMI Damon Runyon Cancer Research Foundation Fellow (DRG-2274-16), who would like to thank S. Montanes-Ordovas for encouraging him to work on human allergic disease.

## Author Contributions

J.O.M., D.F.D., T.M.L., J.A.B., N.A.B., and A.K.S., designed the study. N.B., performed surgeries. J.O.M., D.F.D., C.D., K.M.B., E.Y., collected patient samples and performed single-cell experiments. M.H.W., and T.K.H., provided Seq-Well platform and expertise. H.R.K., performed histologic analyses. J.O.M., D.F.D., S.K.N., and S.W.K., analyzed data. J.O.M., D.F.D., S.K.N., N.A.B., and A.K.S., interpreted data. J.O.M., D.F.D., N.A.B., and A.K.S., wrote the manuscript, with input from all authors.

## Author Information

The authors declare no competing financial interests. Readers are welcome to comment on the online version of the paper. Raw data will be made accessible through DBGap, and partially analyzed data (e.g. **Supplementary Table 2** cells x genes matrix and associated **R script**) will be available upon peer-review and formal publication. **Supplementary Tables 1, 3, 4, and 5** mentioned in the manuscript are currently available upon request. Correspondence and requests for materials should be addressed to N.A.B. and/or A.K.S.

---

**Supplementary Figure 1.**
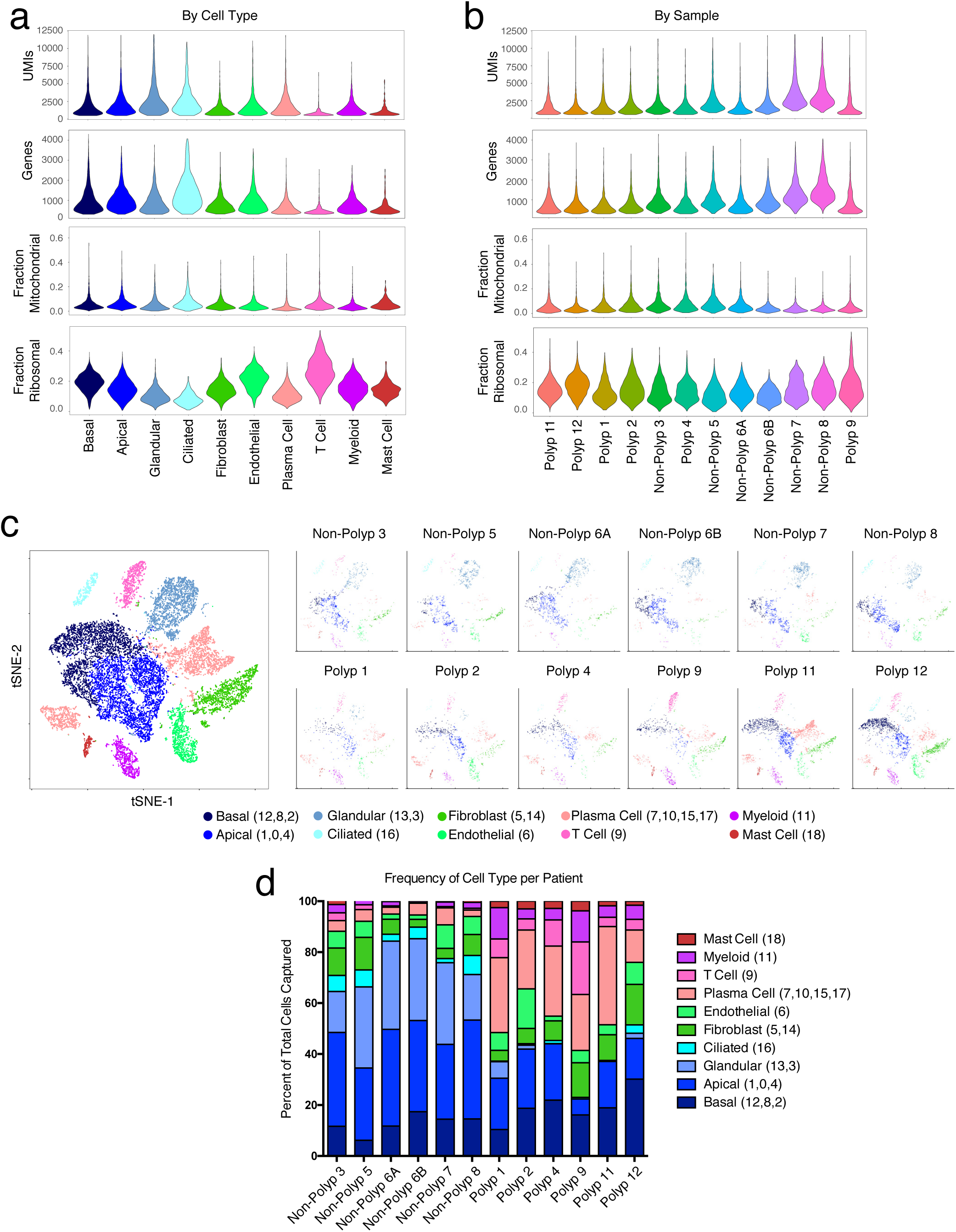
Consistency of cell capture and identification in scRNA-seq patient cohort. **(a)** Number of unique molecular identifiers (nUMI) and genes identified, and fraction of reads mapping to mitochondrial or ribosomal genes across recovered cell types. **(b)** Number of unique molecular identifiers (nUMI) and genes identified, and fraction of reads mapping to mitochondrial or ribosomal genes across patient samples. **(c)** tSNE plot as in **Fig. 1b** colored by cell types across all patients and then separated by sample. **(d)** The percentage of each cell type recovered within each sample.

**Supplementary Figure 2.**
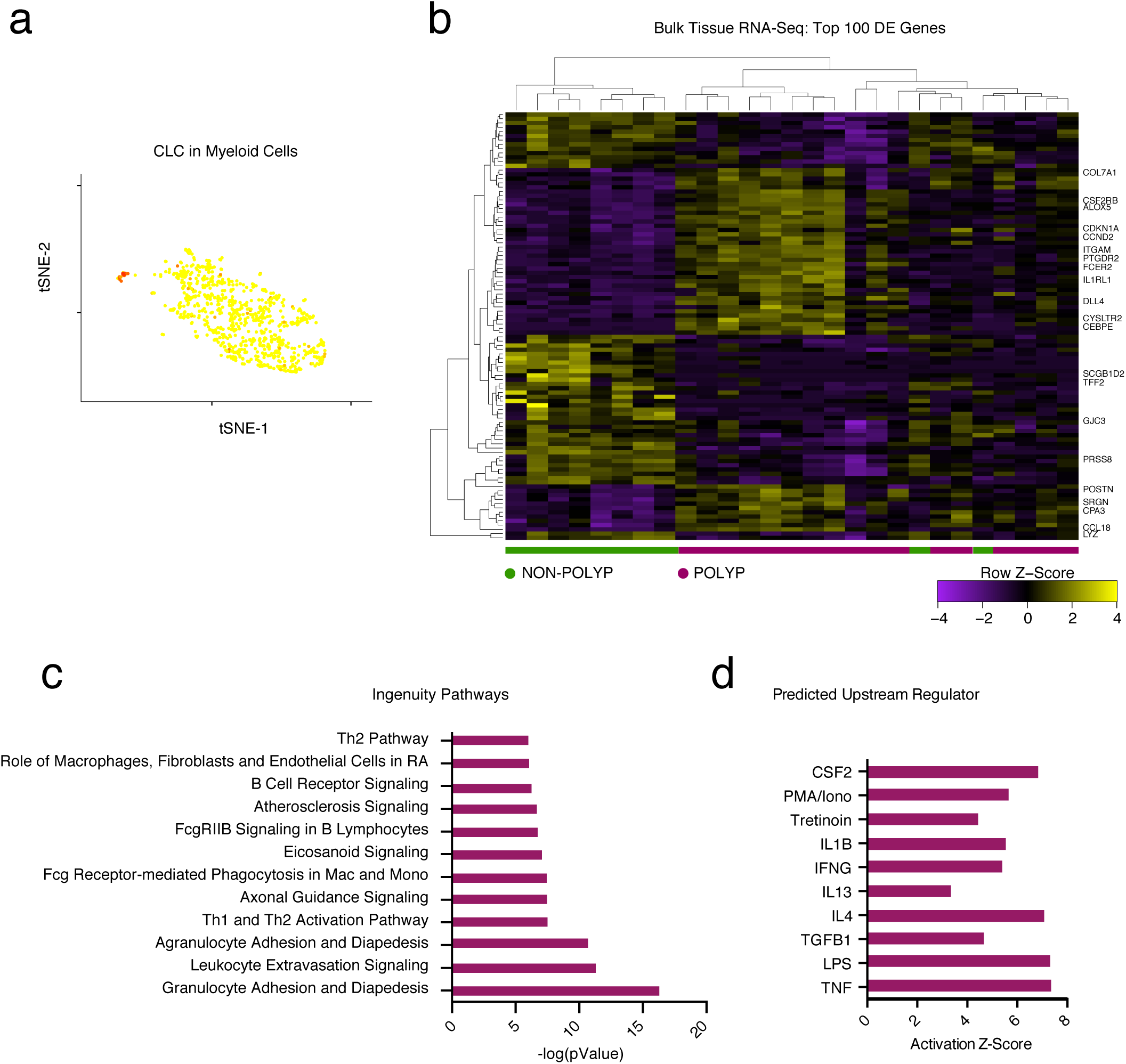
Bulk tissue RNA-seq recovers expected type 2 immune and eosinophilic modules. **(a)** An overlay of *CLC* displaying binned count-based expression level (log(scaled UMI+1) amongst myeloid cells (a pathognomonic gene for eosinophils). **(b)** A row-normalized and row- and column-clustered heatmap over the top 100 positively and negatively differentially-expressed genes (50 in each direction) in bulk tissue RNA-seq of 27 samples from non-polyp (n=9) and polyp (n=18) tissue with select genes displayed; all p<9.03x10-5 for genes displayed corrected for multiple comparisons by Benjamini procedure, (see **Supplementary Table 3** for full gene list and associated statistics). **(c)** The top differentially regulated pathways identified by Ingenuity Pathway Analysis (see Methods) over the top 1,000 differentially expressed genes, as determined by p<0.05 corrected for multiple comparisons by Benjamini procedure, across polyp and non-polyp tissue. **(d)** The predicted upstream regulators based on differentially expressed gene modules in polyp tissue relative to non-polyp determined using Ingenuity Pathway Analysis (see **Methods**).

**Supplementary Figure 3.**
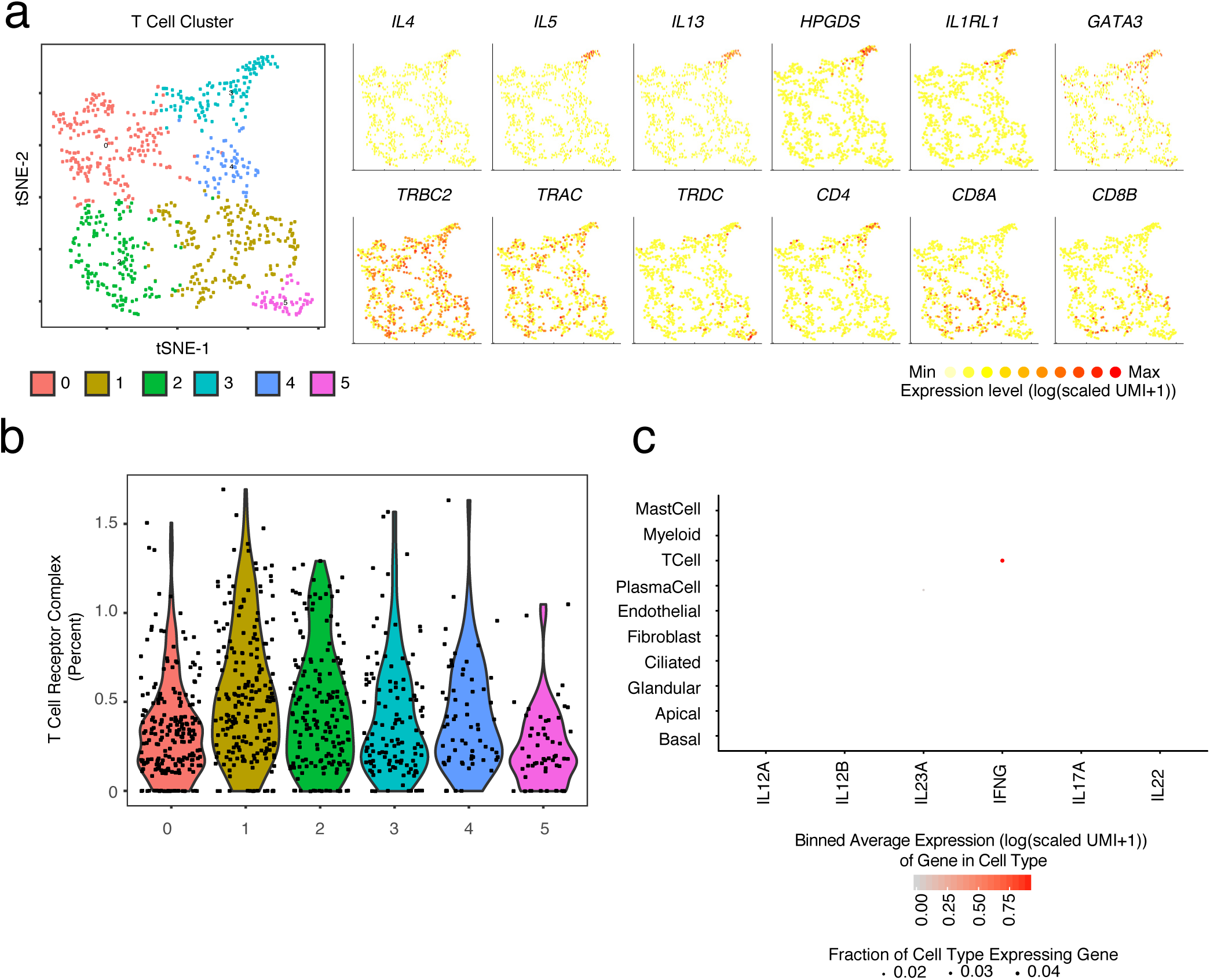
The identities of T cells in type 2 immunity. **(a)** tSNE plot of re-clustered T cells with select gene overlays displaying binned count-based expression level (log(scaled UMI+1) for Th2A-specific genes (top row) and canonical T cell markers (bottom row). **(b)** Violin plot of five identified T cell clusters scored for expression of T cell receptor complex genes (e.g. *TRAC* and *CD3E*, see **Methods**, **Supplementary Table 4**). **(c)** Dot plot of inducers and effectors of Type 1 immunity across all cell types (Note: *IL17F* not detected).

**Supplementary Figure 4.**
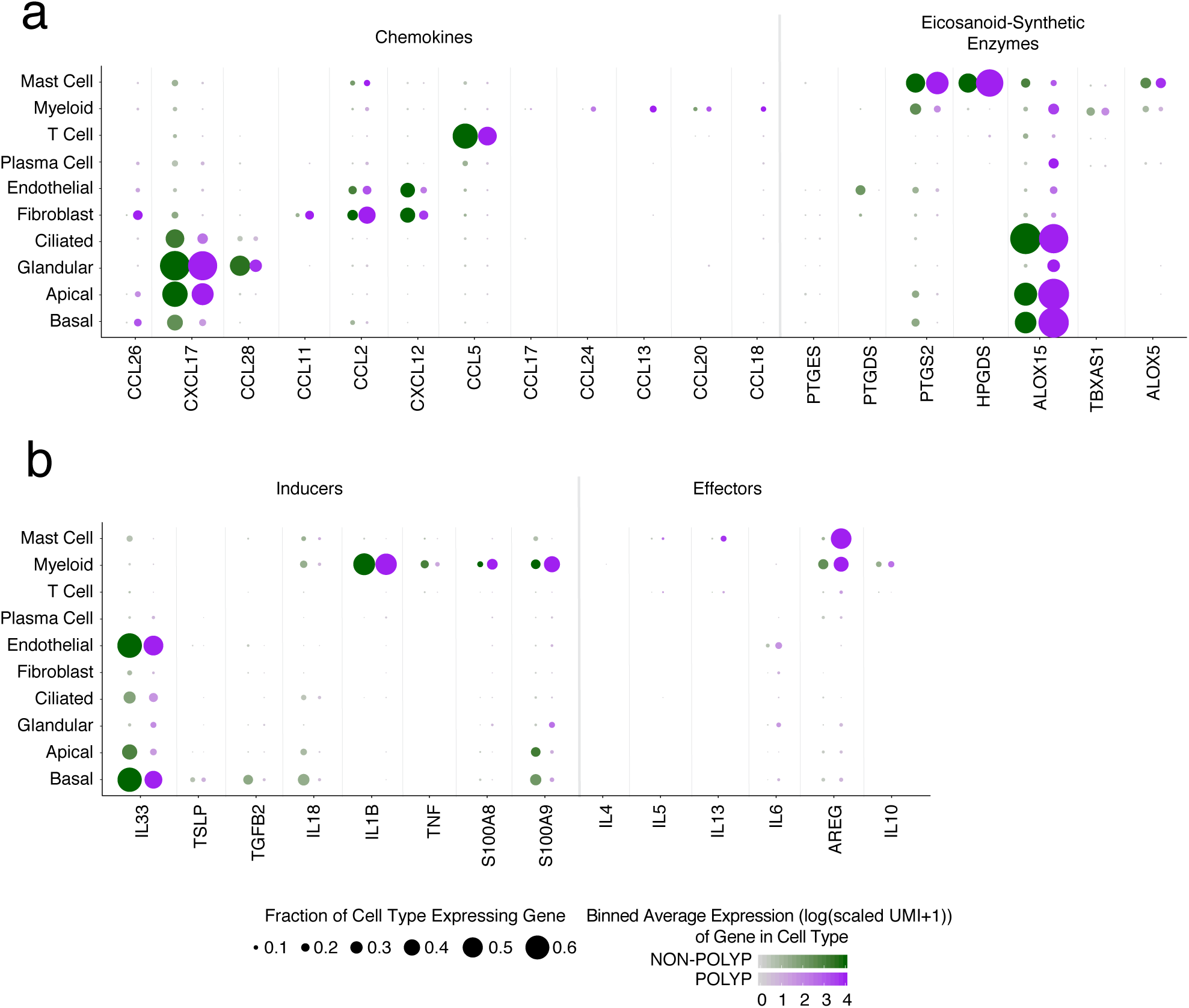
Mapping type 2 inflammatory mediators within non-polyp or polyp ecosystems. **(a)** Dot plots of chemokines and lipid mediators with known roles in type 2 immunity mapped onto cell types divided by non-polyp or polyp disease state, dot size represents fraction of cells within that type expressing, and color intensity binned (log(scaled UMI+1)) gene expression amongst expressing cells (related to **Figure 1d**). **(b)** Dot plot of inducers and effectors of type 2 immunity mapped onto cell types divided by non-polyp or polyp disease state, dot size represents fraction of cells within that type expressing, and color intensity binned (log(scaled UMI+1)) gene expression amongst expressing cells (related to **Figure 1e**).

**Supplementary Figure 5.**
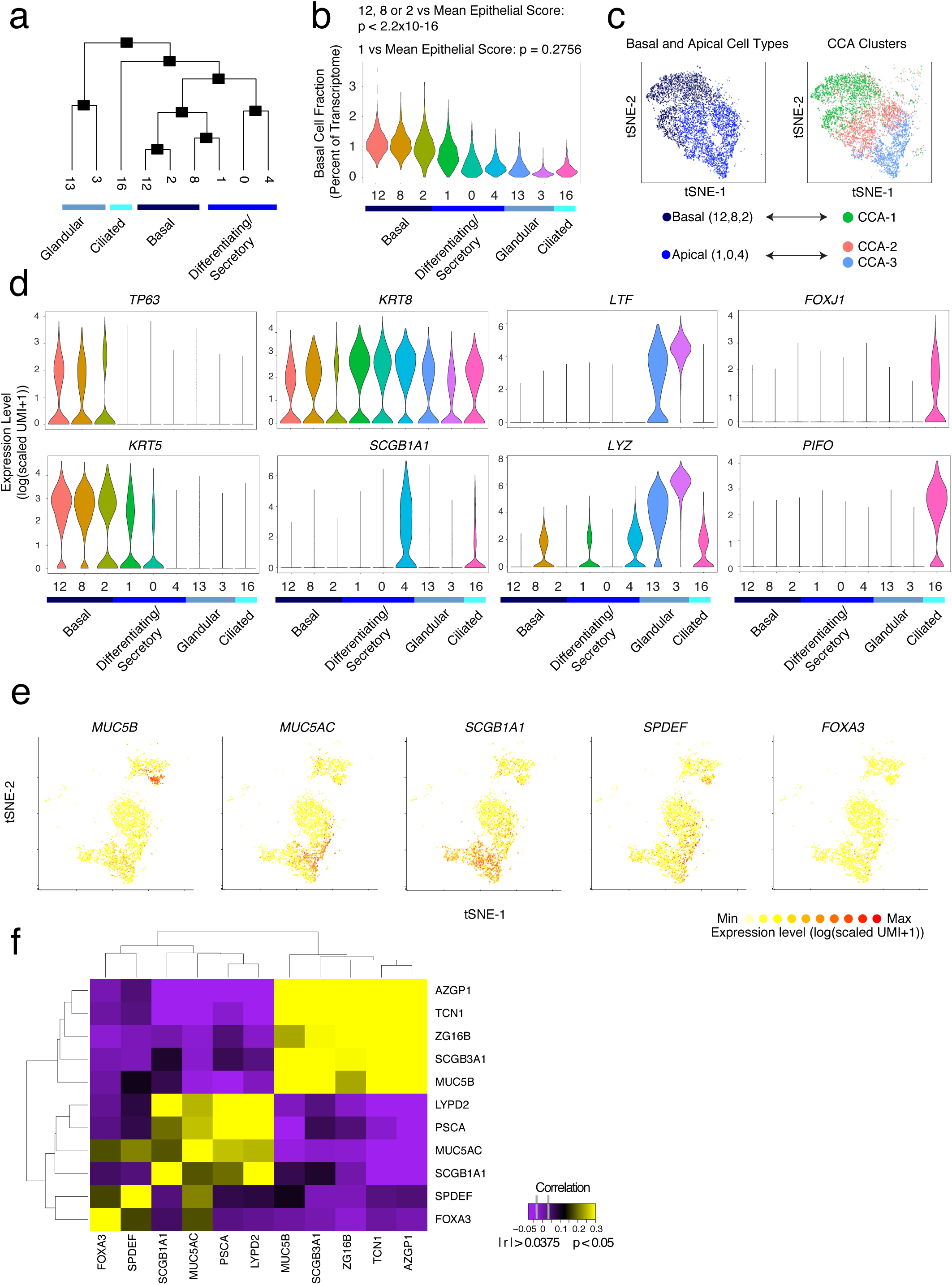
Relationship of epithelial cell clusters and secretory/glandular distinctions. **(a)** A phylogenetic tree based on the average cell from each cluster of epithelial cell clusters in gene-space. **(b)** Violin plot of expression contribution to a cell’s transcriptome of basal cell genes (see **Methods** and **Supplementary Table 4**) across all epithelial cells; *t-test each cluster score vs. the average score of all epithelial cells. **(c)** Canonical correlation analysis (CCA) displaying our cell type annotations for basal and apical cells derived through clustering and biological curation alongside CCA clusters in tSNE space. **(d)** Violin plots for the count-based expression level (log(scaled UMI+1)) of selected marker genes for each identified epithelial cell subset. **(e)** Select overlays on clusters 0 and 4 (differentiating/secretory) and 3 (glandular) displaying binned count-based expression level (log(scaled UMI+1) in tSNE space for canonical goblet (*MUC5B, MUC5AC, SPDEF, FOXA3*) and secretory (*SCGB1A1*) genes. **(f)** A clustered correlation matrix of glandular, goblet, and secretory cell genes; abs(r)> 0.038 is p<0.05 significant based on asymptotic p-values.

**Supplementary Figure 6.**
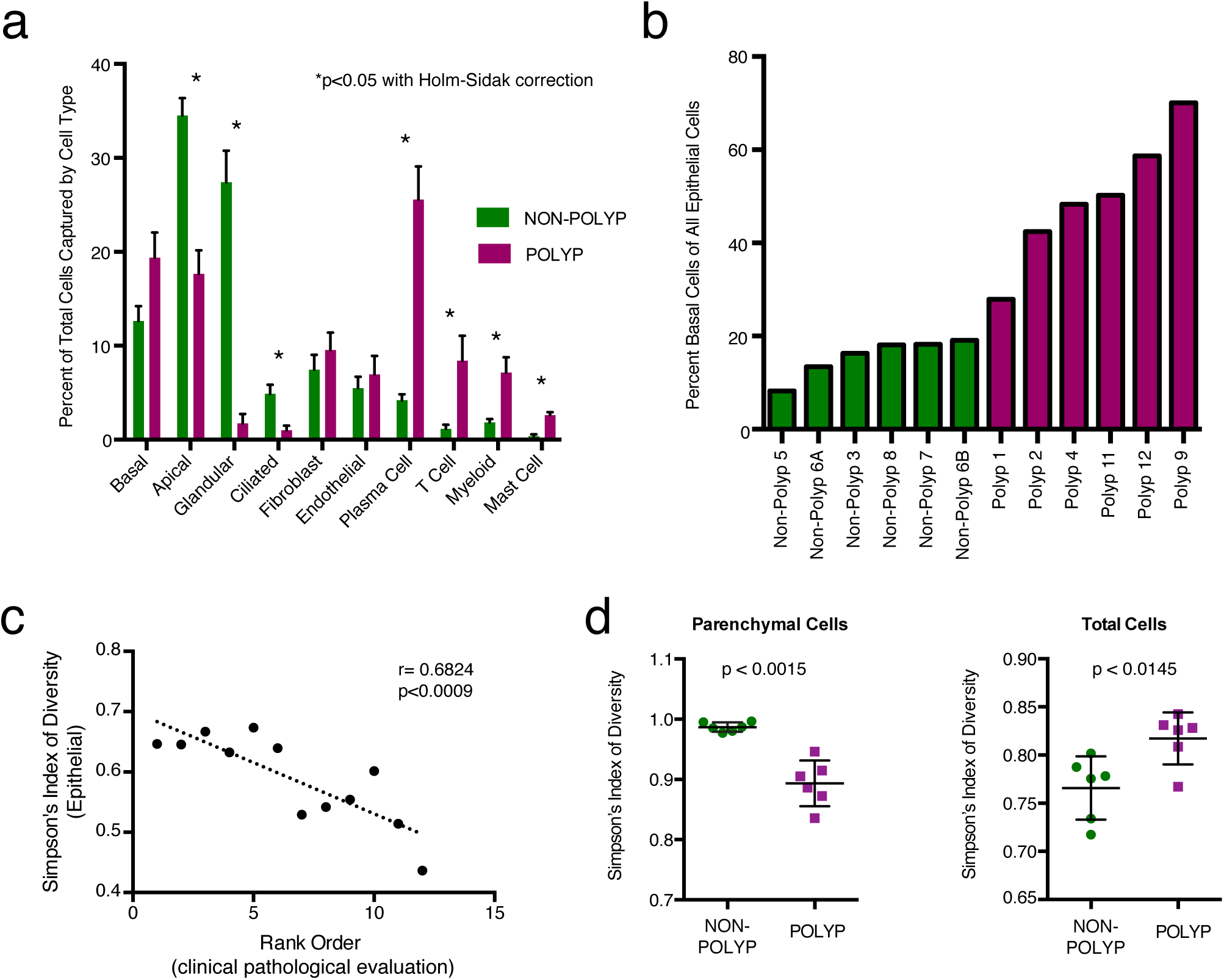
Changes in cellular composition between non-polyp and polyp tissue. **(a)** The frequency of each cell type recovered amongst all cells within each patient sample (n=6 non-polyp, 6 polyp) grouped by disease state; *t-test p<0.05 for Apical, Glandular, Ciliated, Plasma Cell, T Cell, Myeloid and Mast Cell with Holm-Sidak correction for multiple comparisons. **(b)** The frequency of basal cells amongst epithelial cells captured in scRNA-seq data displayed for each sample and colored by non-polyp or polyp designation. **(c)** Correlation of Simpson’s index of diversity calculated over epithelial cells against the ranked order of samples based on clinical pathological evaluation; r=0.6824, p<0.009. **(d)** Simpson’s index of diversity over parenchymal cell types and total cells, an indication of the total richness present within an ecosystem, calculated for each sample; points represent individual samples, *t-test p<0.0015 parenchymal cells, p<0.0145 total cells, non-polp vs. polyp.

**Supplementary Figure 7.**
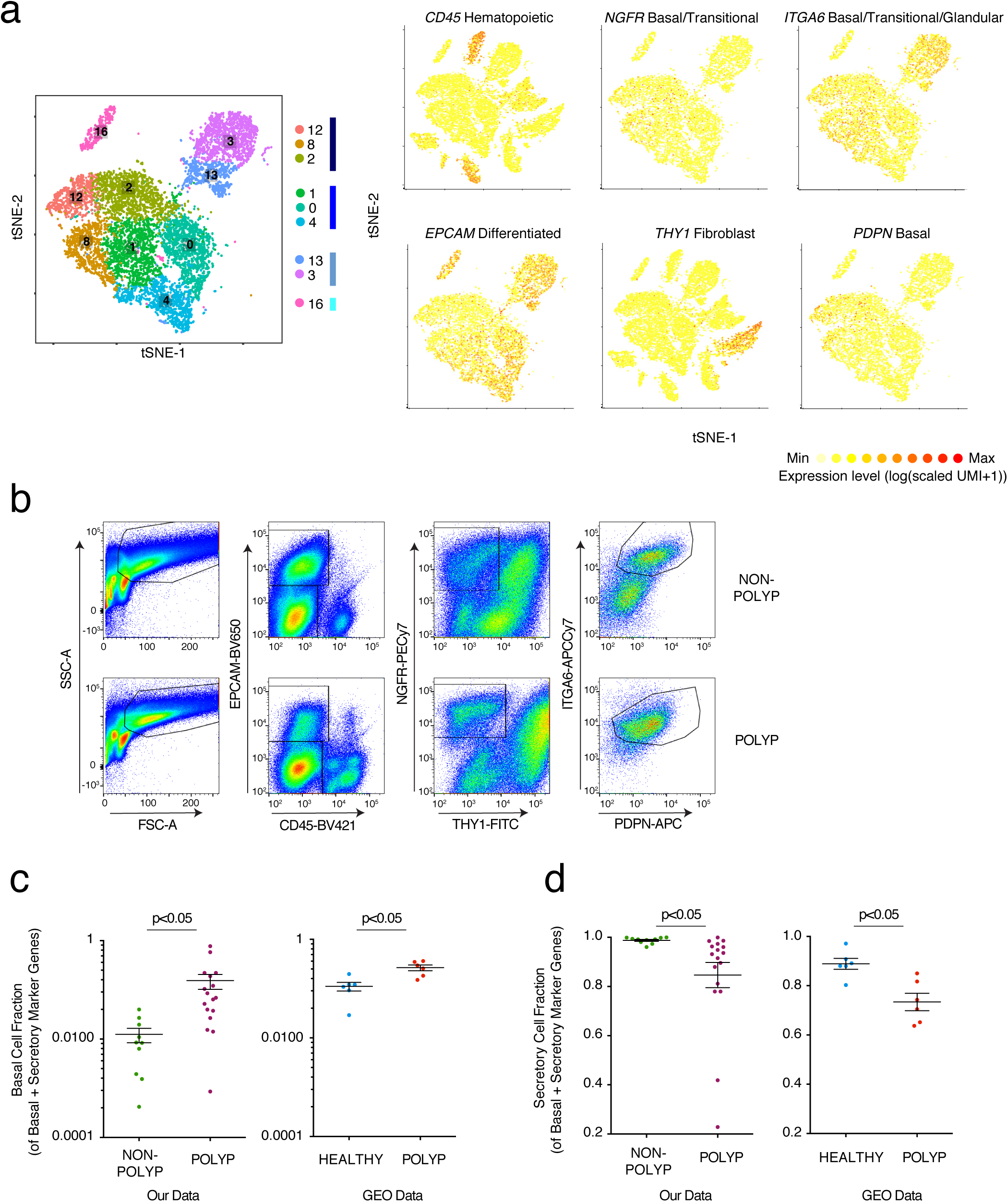
Flow cytometric gating strategy for quantification and isolation of basal cells and validation of basal cell hyperplasia relative to healthy tissue. **(a)** Reproduced from **Figure 2a**: tSNE plot of 10,274 epithelial cells, colored by clusters identified through SNN, with adjacent color bars representing related cell clusters, and overlays displaying binned count-based expression level (log(scaled UMI+1) of selected genes used to negatively (*CD45, EPCAM, THY1*) and positively (*NGFR, ITGA6, PDPN*) identify basal cells. **(b)** Full gating strategy for quantification and isolation of basal cells from non-polyp and polyp tissue, (related to **Fig. 3c,d**). **(c)** Basal cell fraction of transcripts from bulk tissue RNA-seq data of our own data set (related to **Fig 3g,h**) and two GEO data sets (NB: analysis is done per sample and as such no comparisons across the data sets are made) containing healthy and healthy/polyp nasal mucosa biopsies; *t-test p<0.05. **(d)** Secretory cell fraction of transcripts from bulk tissue RNA-seq data of our own data set (related to **Fig 3g,h)** and two GEO data sets (NB: analysis is done per sample and as such no comparisons across the data sets are made) containing healthy and healthy/polyp nasal mucosa biopsies; *t-test p<0.05.

**Supplementary Figure 8.**
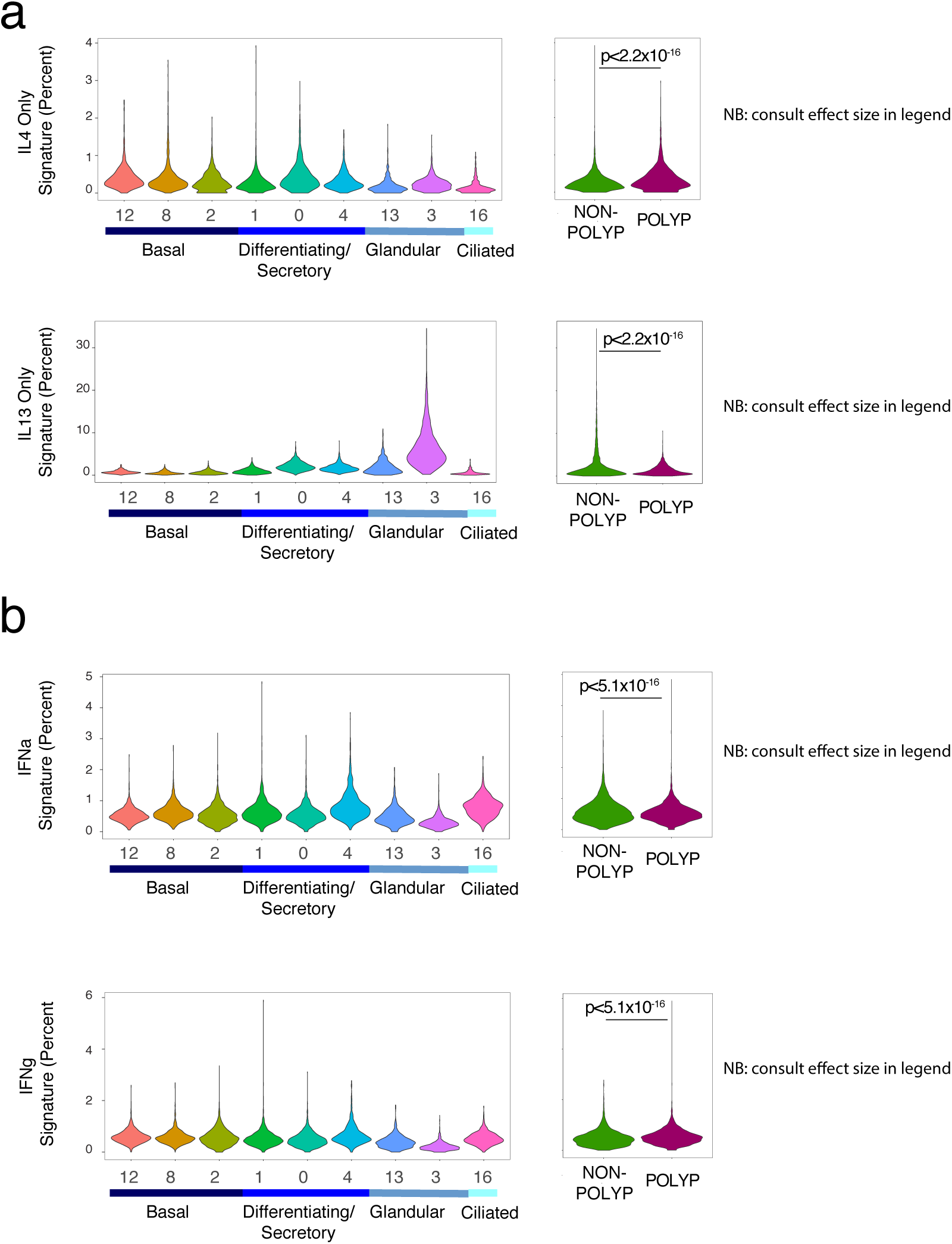
Epithelial cytokine signatures demonstrate Type 2 inflammatory pattern. **(a)** Violin plots of IL-4 or IL-13 uniquely induced gene signatures in respiratory epithelial cell clusters or grouped by disease state presented as expression contribution to a cell’s transcriptome (see **Methods**, **Figure 4b** for shared genes, and **Supplementary Table 4**); *t-test p<2.2x10^−16^, 0.305 effect size IL4 polyp vs. non-polyp and −0.448 effect size IL13 polyp vs non-polyp. **(b)** Violin plots of IFNα or IFNγ induced gene signatures in respiratory epithelial cell clusters or grouped by disease state presented as expression contribution to a cell’s transcriptome (see **Methods**, and **Supplementary Table 4**); *t-test p<5.1x10^−16^ for both, −0.156 effect size IFNα polyp vs. non-polyp and 0.161 effect size IFNγ polyp vs non-polyp.

**Supplementary Figure 9.**
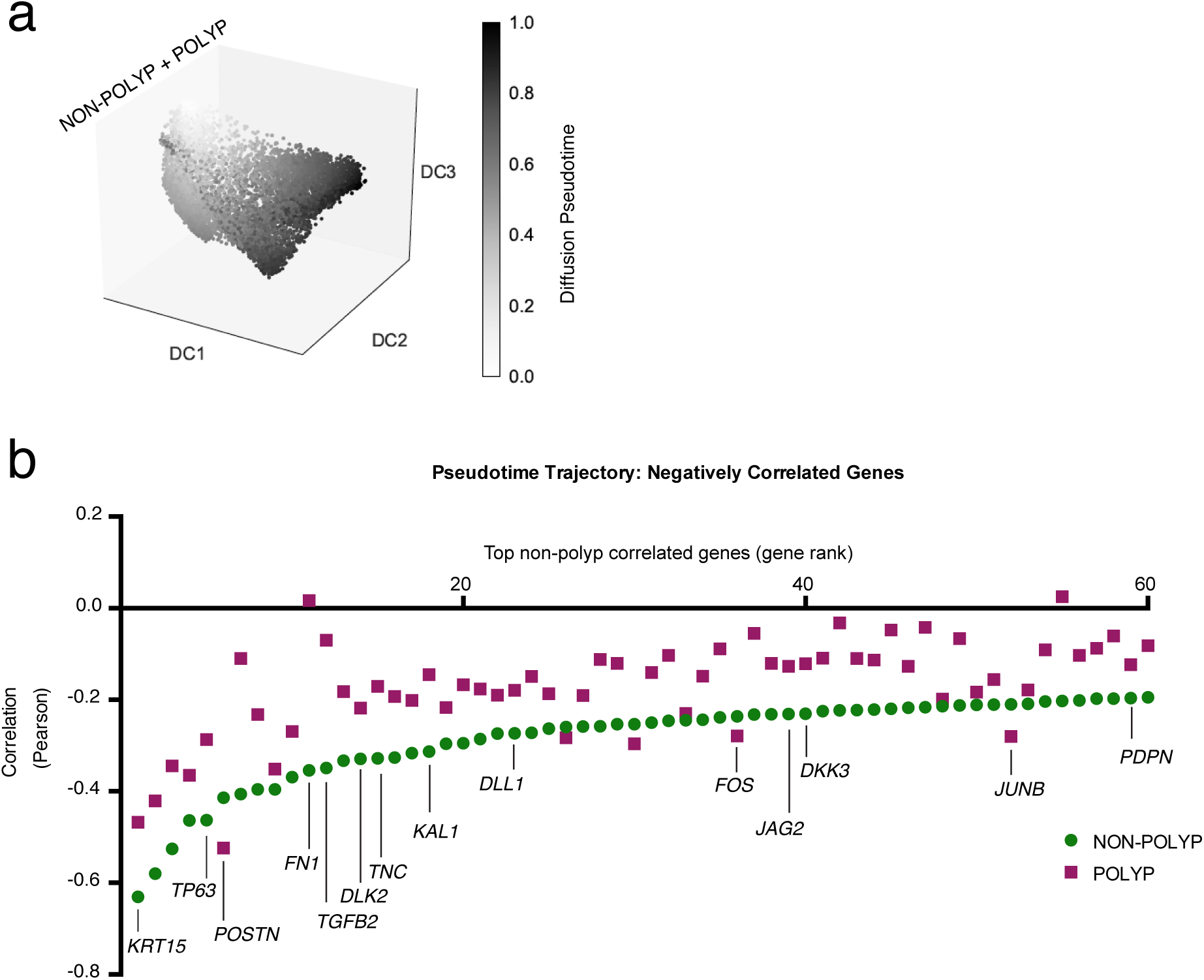
Pseudotime analysis on basal and differentiating/secretory cell clusters. **(a)** Pseudotime analysis using diffusion mapping (see **Methods**) of selected clusters of epithelial cells, here displaying diffusion pseudotime (related to **Fig. 4f**); n= 3,516 cells (clusters 8/1/4), n= 4,064 cells (clusters 12/2/0), and n= 6 non-polyp, 6 polyp samples, diffusion map and DC (diffusion coefficients) are calculated over the set of basal and apical marker genes identified in **Fig. 1a**, see **Supplementary Table 3.** **(b)** The top 60 negatively correlated genes expressed in non-polyp cells with pseudotime trajectory and Pearson correlation values for genes in polyp cells also displayed; differential correlation coefficient analysis using Fisher’s Z-statistic, accounting for number of cells in each group (specific genes highlighted all > 2 Z, full results including Bonferroni corrected p values in **Supplementary Table 3**).

**Supplementary Figure 10.**
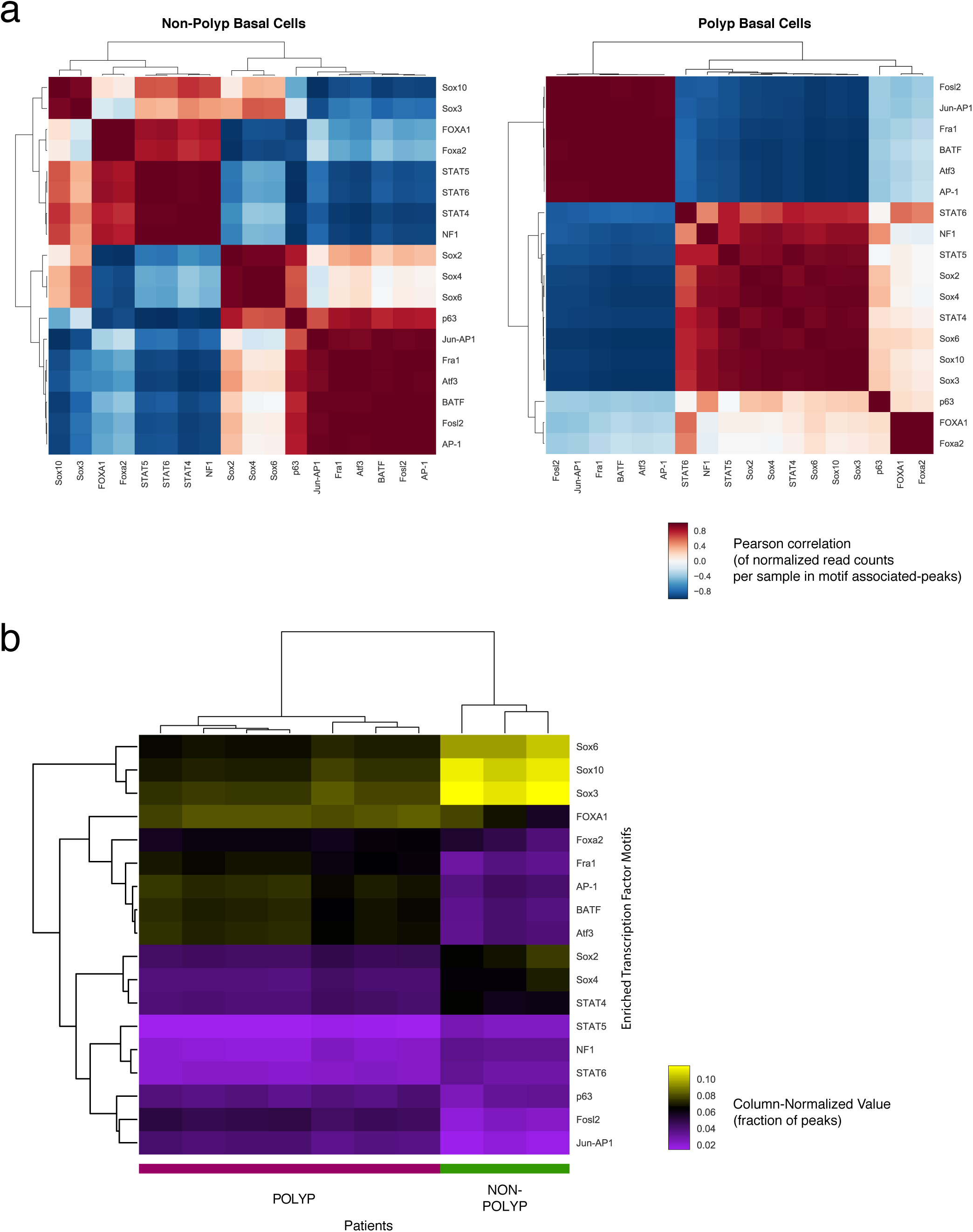
Transcriptional motif enrichments in non-polyp and polyp basal cells. **(a)** Correlation matrices (row and column clustered) of the normalized read counts per sample in motif associated-peaks for non-polyp or polyp samples; Pearson correlation, n=3 non-polyp, n=7 polyp. **(b)** A column-normalized heatmap (row and column clustered) for the fraction of peaks with a motif corresponding to accessibility of the respective transcription factor displayed by patient; n=3 non-polyp, n=7 polyp.

